# Membrane-cytoskeleton mechanical feedback mediated by myosin-I controls phagocytic efficiency

**DOI:** 10.1101/433631

**Authors:** Sarah R. Barger, Nicholas S. Reilly, Maria S. Shutova, Qingsen Li, Paolo Maiuri, Mark S. Mooseker, Richard A. Flavell, Tatyana Svitkina, Patrick W. Oakes, Mira Krendel, Nils Gauthier

**Affiliations:** Cell and Developmental Biology Department, State University of New York Upstate Medical University, Syracuse, NY; Department of Physics, University of Rochester, Rochester, NY; Department of Biology, University of Pennsylvania, Philadelphia, PA; IFOM, FIRC Institute of Molecular Oncology, Milan, Italy; Molecular, Cellular and Developmental Biology, Yale University, New Haven, CT; Department of Immunobiology, Yale University School of Medicine, New Haven, CT; Howard Hughes Medical Institute, Yale University, New Haven, CT; Department of Biology, University of Rochester, Rochester, NY; Mechanobiology Institute, MBI, National University of Singapore, Singapore

**Keywords:** macrophage, phagocytosis, actin, myosin, membrane tension

## Abstract

Phagocytosis of invading pathogens or cellular debris requires a dramatic change in cell shape driven by actin polymerization. For antibody-covered targets, phagocytosis is thought to proceed through the sequential engagement of Fc-receptors on the phagocyte with antibodies on the target surface, leading to the extension and closure of the phagocytic cup around the target. We have found that two actin-dependent molecular motors, class 1 myosins myosin 1e and myosin 1f, are specifically localized to Fc-receptor adhesions and required for efficient phagocytosis of antibody-opsonized targets. Using primary macrophages lacking both myosin 1e and myosin 1f, we found that without the actin-membrane linkage mediated by these myosins, the organization of individual adhesions is compromised, leading to excessive actin polymerization, slower adhesion turnover, and deficient phagocytic internalization. This work identifies a novel role for class 1 myosins in coordinated adhesion turnover during phagocytosis and supports a model for a membrane-tension based feedback mechanism for phagocytic cup closure.

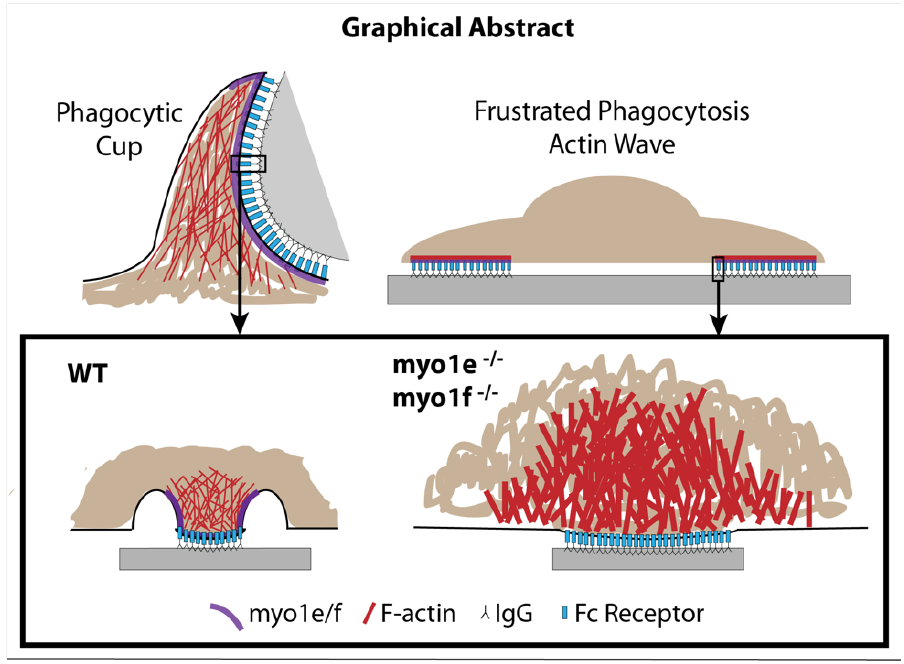

## Introduction

Phagocytosis is a critical immune response that requires coordinated adhesion, membrane rearrangement, and dynamic remodeling of the actin cytoskeleton^1^. Internalization via Fcg receptors (FcRs), which bind the conserved domain of immunoglobulins, involves several stages, beginning with the clustering of FcRs that activate downstream signaling pathways to induce assembly of an actin-rich, cup-like structure (the phagocytic cup) that surrounds the target^2^. The plasma membrane of the phagocytic cup is extended by the force of branched actin polymerization and, if a target is particularly large, additional membrane from intracellular stores is added to the cup by exocytosis^3^. Cup fusion results in a *de novo* membrane-bound organelle (the phagosome), which is shuttled further into the cell for processing and degradation^4^. While the signaling pathways that link FcR clustering to the initiation of F-actin assembly are well understood^5^, extension and closure of the phagocytic cup, which requires regulated actin polymerization and coactive membrane deformation, remains enigmatic.

A number of past studies have revealed phagocytosis to be an intricate biophysical process^6^. For a successful phagocytic event, the force of actin polymerization within the extending arms of the phagocytic cup must overcome mechanical properties of the cell itself, namely membrane and cortical tension. However, as a phagocyte ingests a target, both membrane and cortical tension increase^7–9^, and these properties in turn can regulate addition of new membrane through exocytosis. Over the course of internalization, macrophages experience a steep increase in membrane tension, which triggers exocytosis of intracellular membrane stores that increase cell surface area for internalization^9^. Whether actin polymerization is regulated biophysically to successfully close the phagocytic cup has yet to be explored.

Here, we report that two class 1 myosins, myosin 1e (myo1e) and myosin 1f (myo1f), are regulating actin dynamics by a novel mechanism to allow efficient engulfment of phagocytic targets. Following initial target binding, F-actin is assembled at discrete FcR adhesions between the phagocyte and the IgG-coated particle, and subsequent cup extension is believed to be driven by the formation of additional Fc receptor-IgG bonds in a zipper-like fashion along the target^10^. We have discovered that myo1e/f, small monomeric actin-based motors that can bind to the actin cytoskeleton through their motor domains and the plasma membrane through their tails, are associated with Fc-receptor adhesions and control membrane tension at these sites throughout phagocytosis. Using a myo1e/f double knockout (dKO) mouse model, we found that macrophages lacking these myosins assemble phagocytic cups of clumped and disorganized actin, exhibit slower FcR adhesion turnover and, as a result, are deficient at internalizing targets. By locally organizing membrane around FcR adhesion sites, myo1e/f work to spatially confine FcR signaling. Overall, this work provides an unexpected biophysical explanation for how actin dynamics are precisely controlled to promote extension and closure of the phagocytic cup.

## Results

### Myo1e and myo1f are enriched at the actin-membrane interface of the phagocytic cup and are required for efficient phagocytosis

We used fluorescence microscopy to examine myo1e and myo1f localization both in fixed cells and throughout the time-course of phagocytosis in live cells. RAW264.7 (RAW) macrophages transfected with fluorescently tagged myo1e or myo1f and actin-labeling constructs were challenged to engulf 6 μm latex beads opsonized in mouse IgG. We found that during bead ingestion, myo1e was recruited to the cup and co-localized with the extending belt of phagocytic actin, as has been previously observed^11^, yet slightly preceded actin at the leading edge of the cup (arrowheads) (Fig. 1a-c, Supplementary Fig. 1d, Supplementary Movie 1). Myo1f exhibited similar behavior during engulfment (Supplementary Fig. 1a-c, Fig. 1d) and, when co-expressed, the two myosins showed near perfect colocalization at the cup tip (Fig. 1e). These observations were reinforced by immunostaining of endogenous myo1e and actin during phagocytosis in primary murine bone-marrow derived macrophages (BMDM) (Fig. 1f). These results indicate a clear recruitment and enrichment of myo1e/f at the progressing cup during phagocytosis, with a strong colocalization with the actin belt, as well as a unique localization at the extreme leading edge of the cup, particularly evident near the time of cup closure (Fig. 1d and Supplementary Fig. 1d, arrowheads).

**Fig. 1:**
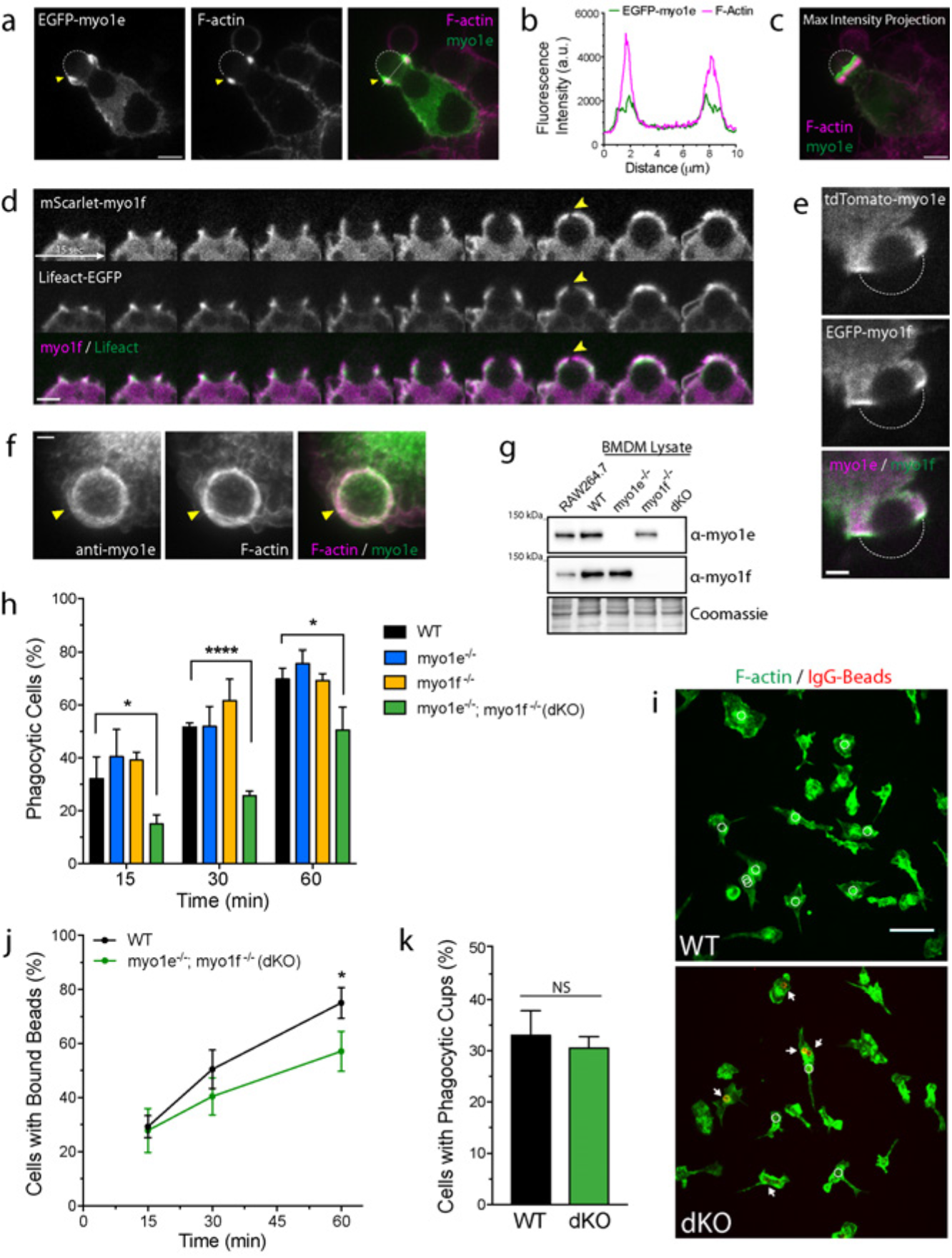
Myo1e and myo1f are enriched at the actin-membrane interface of the phagocytic cup and are required for efficient phagocytosis. a) Representative confocal section of a RAW264.7 macrophage transfected with EGFP-myo1e engulfing a 6 μm IgG-coated bead and stained with fluorescently-labeled phalloidin. Yellow arrowhead points to the phagocytic cup, and the bead is outlined by a dotted white line. Scale bar, 5 μm. See also Supplementary Movie 1. b) Line scan of EGFPmyo1e and F-actin intensity along the white line in (a) Merge to illustrate actin and myo1e colocalization at the phagocytic cup. c) Maximum intensity projections of (a) Merge show that myo1e precedes actin at the leading edge of the phagocytic cup. d) Time-lapse montage of RAW macrophage expressing mScarlet-myo1f and Lifeact-EGFP engulfing 8 μm IgG-coated bead. Yellow arrowhead points to myo1f preceding Factin, particularly at cup closure. Scale bar, 5 μm. e) Myo1e and myo1f colocalize at the edge of phagocytic cup. Representative confocal section of RAW macrophage cotransfected with tdTomato-myo1e and EGFP-myo1f engulfing a 6 μm IgG-coated bead (dotted line). Scale bar, 2 μm. f) Endogenous myo1e colocalizes with F-actin at the phagocytic cup (yellow arrow, bead not outlined). WT BMDM was challenged with 6 μm IgG-coated beads, fixed and stained with myo1e antibody and Alexa-Fluor-488-phalloidin. Total intensity projection of a confocal z-stack. Scale bar, 2 μm. g) Western blots of myo1e/f in RAW264.7 murine macrophage cell line and bone-marrow derived macrophages (BMDM) of WT (myo1e^+/+^; myo1f^+/+^), myo1e-/-, myo1f^-/-^, and myo1e^-/-^; myo1f^-/-^ double knockout (dKO) mice. Equal protein loading was verified by Coomassie Blue staining. h) Graph of percentage (mean ± SEM) of phagocytic cells that successfully internalized at least one bead. WT, myo1e^-/-^, myo1f^-/-^ and myo1e^-/-^ and myo1f^-/-^ (dKO) BMDM were challenged with 6 μm IgG-coated latex beads and analyzed at 15, 30, 60 minutes. Data from 3 independent experiments (*= p<0.05 ****= p<0.0001). i) Representative images from phagocytosis assay of WT and dKO macrophages at 15-minute time point. Cells are stained with phalloidin (green) and un-internalized beads are stained red. Unfilled white circles denote internalized beads (not visible in the fluorescence image), while white arrows point to uninternalized beads exposed to stain. Scale bar, 50 μm. j) Percentage of cells (mean ± SD) that bound at least one bead during phagocytosis time course (*= p<0.05). k) Percentage of cells (mean ± SD) that formed actin-rich phagocytic cups at 10-minute time point (p = 0.44).

After establishing a clear colocalization between actin and myo1e/f during phagocytosis, we next investigated the relationship between these myosins and plasma membrane phospholipids. During phagocytic internalization, lipid composition within the cup undergoes a series of changes, which parallel and likely regulate sequential stages in actin assembly and dynamics^12^. PtdIns(4,5)P_2_ (PIP2) populates the extending arms of the phagocytic cup and is replaced by PtdIns(3,4,5)P_3_ (PIP3) as the cup closes^13,14^. PtdIns(3)P is then a major lipid during phagosome maturation^15^. As myo1e and myo1f contain putative Pleckstrin Homology (PH) domains, capable of interacting with both PIP2 and PIP3^16, 17^, we tested whether the two myosins occupied the same regions as these lipids. RAW macrophages were co-transfected with fluorescent myo1e/f and the EGFP-tagged lipid sensors PLCδ-PH, for PtdIns(4,5)P_2_, and AKT-PH, for both PtdIns(3,4,5)P_3_ and PtdIns(3,4)P_2_ (Supplementary Fig. 2a)^13,18^. As a negative control, we used PKCd-C1, a sensor for diacylglycerol^19^, which is known to be most enriched in sealed phagosomes^20^. By assessing the percentage of cells that displayed myosin and lipid sensor colocalization, we concluded that myo1e and myo1f colocalized predominantly with PtdIns(3,4,5)P_3_ at the phagocytic cup (Supplementary Fig. 2b). Colocalization with PtdIns(4,5)P_2_ was also observed, but at a lower frequency. To test the dependence of myosin localization at the cup on phospholipid accumulation, we assessed cup recruitment of myo1e and myo1f following treatment with LY294002, an inhibitor of PI3K, thus blocking PtdIns(3,4,5)P_3_ production at the phagocytic cup^21,22^. Both myo1e and myo1f still localized to the phagocytic cup, although to a lesser extent (Supplemental Fig. 2c-d). Together, these results show that myo1e and myo1f may interact with both cytoskeleton and membrane at their location within the cup through actin and phosphoinositide binding respectively, but do not solely depend on PtdIns(3,4,5)P_3_ for their recruitment.

The observed localization and behavior of the two myosin-Is strongly argue for a role in the coordination of actin dynamics and membrane remodeling at the cup during phagocytosis. In order to test whether myo1e and myo1f were required for phagocytosis, we isolated bone-marrow derived macrophages (BMDM) from wild type mice, myo1e KO mice^23^, and myo1f KO mice^24^. Both KO strains were created using a constitutive (germline) knockout approach, so that cultured macrophages completely lack myo1e or myo1f (Fig. 1g). Given the similarity in myo1e/f domain structure and the likelihood of functional redundancy, as suggested by our localization results, we also generated a double knockout (dKO) mouse to best understand the role of these proteins in phagocytosis. To measure phagocytic activity of BMDMs, we challenged cells to engulf 6 μm IgG-coated latex beads. We identified un-internalized beads by fixing cells and staining beads prior to cell permeabilization (see Supplementary Fig. 1e). By quantifying the percentage of cells that engulfed at least one latex bead, we found that macrophages lacking either myo1e or myo1f performed similar to control cells (Fig. 1h). However, when both myosins were knocked out, macrophages appeared to be engulfing beads more slowly, with a significantly lower percentage of phagocytosing cells at each time point (Fig. 1h-l). This result suggests myo1e and myo1f are required for efficient phagocytosis, yet perform functionally redundant roles, and thus all subsequent experiments with BMDMs were conducted using WT and dKO macrophages.

### Myo1e and myo1f are not required for FcR signaling, target binding, or phagocytic cup formation

With myo1e and myo1f binding both actin filaments and membrane lipids, the lack of these myosins could potentially affect FcR clustering and activity at the plasma membrane or the ability of cells to assemble actin to form the phagocytic cup. To test whether any of these mechanisms could account for the observed phagocytosis defect, we first assessed downstream signaling from FcRs by stimulating BMDM with anti-FcR antibody followed by antibody cross-linking at 37ºC for different periods of time. We found that levels of phosphorylated Syk, Akt, and downstream ERK signaling in dKO cells were indistinguishable from those in WT macrophages (Supplementary Fig. 1f-g). We next tested whether FcR surface presentation or interactions with the antibody-coated targets were affected. Flow cytometry showed that FcR surface expression was unaffected in dKO cells (Supplementary Fig. 1h). In addition, the percentage of cells that were simply associated with a bead, whether external (bound) or internal (ingested), was similar between WT and dKO BMDM (Fig. 1j). Notably, at 15 minutes, when a 50% defect in bead ingestion was observed (Fig. 1h), we detected no statistically significant difference in bead association between the WT and dKO macrophages (Fig. 1j). Both the unchanged receptor density and efficiency of early bead binding suggested that myosin loss did not affect FcR behavior on the cell surface.

We next questioned whether the lack of myo1e/f was affecting phagocytic cup formation. WT and dKO BMDM were exposed to IgG-coated latex beads for 10 minutes, then fixed and stained with fluorescently labeled phalloidin to identify phagocytic cups.

Surprisingly, the percentage of cells with phagocytic cups did not differ between WT and dKO cells (Fig. 1k). Given the successful bead binding and formation of phagocytic cups in dKO macrophages, the lag in complete bead internalization that we observed can only be due to a defect during cup progression and/or closure in the absence of myo1e/f.

### Myo1e and myo1f are not required for contractility during frustrated phagocytosis

As myo1e/f are molecular motors, and contractility is known to be required for the completion of phagocytosis^25^, the inability of dKO cells to efficiently close their phagocytic cups may be the result of reduced contractility applied by the macrophage on the target. Because myo1e has been previously proposed to promote phagosome closure by driving contraction of the phagocytic cup^26^, we set out to test whether macrophages lacking myo1e/f are deficient in contractile force production. We employed traction force microscopy (TFM), which has recently been validated to measure contractility during a 2D frustrated phagocytosis assay^27^, to test the contribution of myo1e/f to contractile force generation. During frustrated phagocytosis, cells spread on an IgG-coated coverslip, which stimulates actin and membrane rearrangement that faithfully mimics the 3D process^28–31^. For TFM, flat but deformable polyacrylamide gels were prepared for frustrated phagocytosis by first coating with BSA followed by anti-BSA mouse antibody. WT and dKO macrophages were then dropped on to the gels and imaged for 30 minutes while spreading (Fig. 2a, Supplementary Movie 2). Over time the mean strain energy produced by the dKO cells did not significantly differ from that of WT macrophages (Fig. 2b-c). Quantification of the maximum strain energy also did not reveal any significant differences in contractile activity that could be attributed to myo1e/f (Fig. 2d). Thus, myo1e/f appear not to contribute to the contractile force applied by macrophages to the target during FcR-mediated phagocytosis, as measured by this assay.

**Fig. 2:**
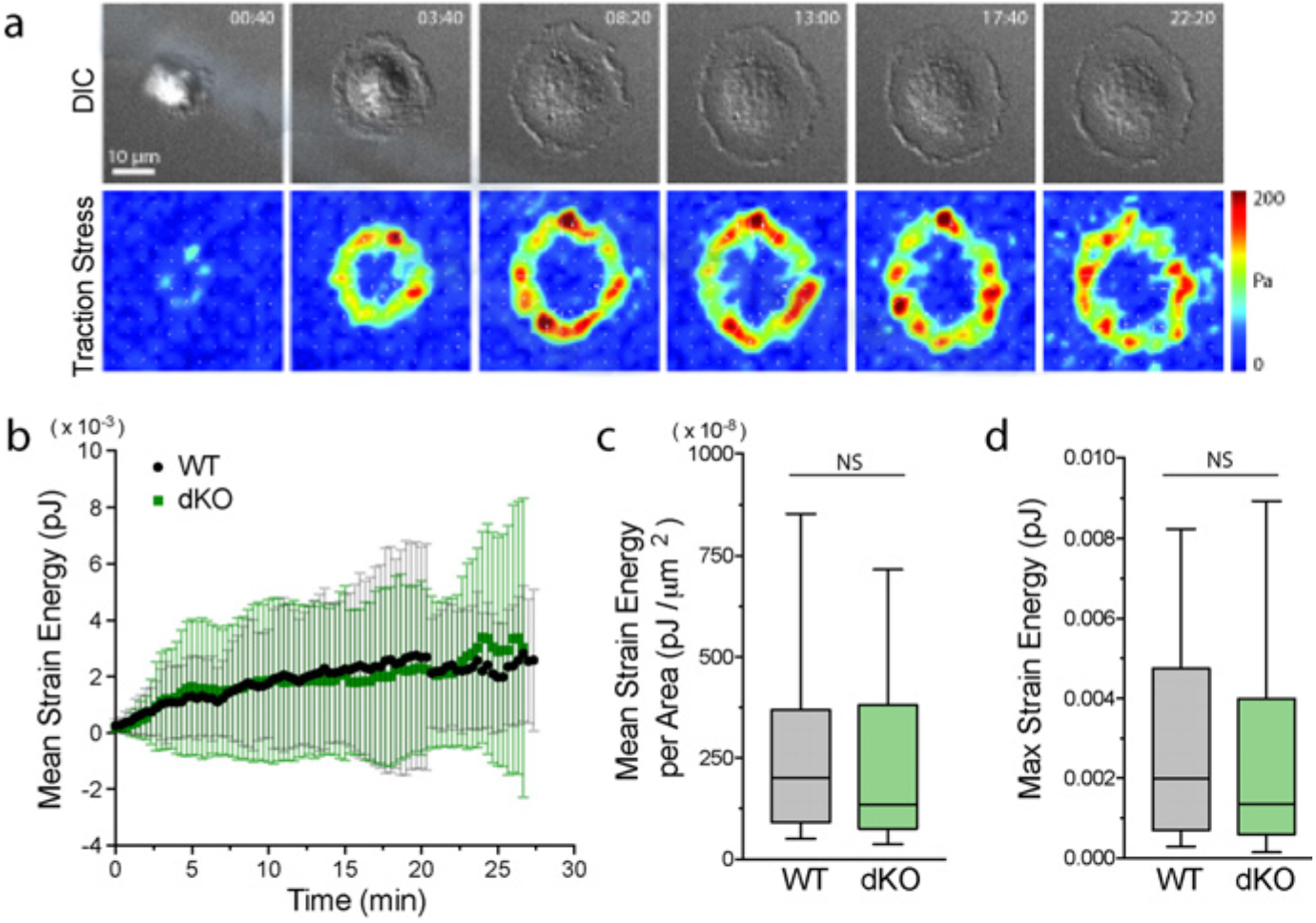
Myole and myolf do not contribute to phagocytic contractility. a) Representative time-lapse montage of BMDM (dKO macrophage) performing frustrated phagocytosis and exhibiting traction forces. Differential interference contrast (DIC) imaging (top) beside traction force map (bottom). The magnitude of the brightness in the traction map corresponds to the magnitude of the stress (i.e. a pixel value of 50 = 50 Pa), with the pixel intensity values color-coded as indicated by the color wedge on the right. Scale bar, 10 μm. See also Supplementary Movie 2. b) Graph of mean strain energy over time during spreading. WT and dKO macrophages performed frustrated phagocytosis for TFM measurements. Data from 2 independent experiments (n= > 40 cells per genotype). c) Graph of mean strain energy per unit of cell area. Box and whisker plot shows median, 25^th^ and 75^th^ percentile, with error bars depicting maximum and minimum data points. Two outliers have been removed (p=0.43). d) Graph of maximum strain energy measured over 30 minutes of cell spreading. Box and whisker plot shows median, 25^th^ and 75^th^ percentile, with error bars depicting maximum and minimum data points. Four outliers (out of > 40 cells) have been removed (p = 0.48).

### Myole and myolf do not participate in focal exocytosis

Focal exocytosis, the local insertion of intracellular membrane at the phagocytic cup, allows cells to fully engulf large targets, such as the 6 μm IgG-coated beads used in our study. Because class 1 myosins are known to localize to endosomes and exocytic vesicles^32–34^, we hypothesized that myo1e/f may be mediating membrane vesicle delivery at the phagocytic cup explaining the slower cup closure observed in dKO BMDM. To test this hypothesis, we challenged cells to engulf smaller targets (2 μm latex beads), which should not require focal exocytosis activity^21^. Despite the smaller target size, we found that myo1e still localized to the phagocytic cup (Supplementary Fig. 3a). This agrees with our finding that the motor proteins are recruited in a PI3K-independent manner, which is known to be dispensable for the ingestion of smaller targets^35^. We found that dKO macrophages still exhibited slower uptake of 2 μm beads, with no differences in bead binding (Supplementary Fig. 3b-c). To further investigate membrane dynamics, we utilized a lipid-based dye, *N*-(3-Triethylammomumpropyl)-4-(4-(dibutylamino) styryl) pyridinium dibromide (FM 1–43), to measure increases in cell surface area, regardless of the specific membrane source added, during frustrated phagocytosis. As we have previously shown using this assay, macrophages undergoing frustrated phagocytosis increase their cell surface area by ∼40%, which is reflected as an increase in FM 1–43 intensity^9^. Using WT and dKO macrophages, we observed no difference in FM 1–43 intensity over the course of cell spreading (Supplementary Fig. 3d-f). In past studies, defects in focal exocytosis and subsequent failure to extend the phagocytic cup have been demonstrated by a reduction in spread cell area on IgG^36–38^. We thus quantified spread cell area during 25 minutes of frustrated phagocytosis and found no difference in dKO cell area compared to WT macrophages over time (Supplementary Fig. 3g). Moreover, the velocity of the leading edge during cell spreading was also measured and showed no difference between WT and dKO cells (Supplementary Fig. 3h). Finally, we used RAW macrophages to appraise myo1e/f localization with respect to focal exocytosis markers, such as VAMP3^39^. We detected no colocalization between myo1e/f and VAMP3, finding that class 1 myosins appear spatially separated from focal exocytosis machinery within the cup (Supplementary Fig. 3i).

Together, these data affirm that myo1e and myo1f are not involved in focal exocytosis.

### Myole and myolf localize to FcR-actin adhesions during frustrated phagocytosis

Having ruled out binding, initial cup formation, and early pseudopod extension, as well as contractile force generation and focal exocytosis, as potential activities that could explain the observed phagocytic defect affecting dKO BMDM, we decided to investigate actin dynamics within the phagocytic cup. As early pseudopod protrusion may not be affected, based on our finding that initial spreading velocity is similar in WT and dKO cells (Supplementary Fig. 3g), we were particularly interested in actin turnover following the initial cup formation. Continued actin polymerization during the final stages of phagocytosis is required to push the membrane forward to complete internalization. To better visualize phagosomal actin, we imaged cells performing frustrated phagocytosis by total internal reflection fluorescence microscopy (TIRFM). While performing frustrated phagocytosis, macrophages are stimulated to form dynamic actin waves, cleared of cortical actin in their centers, that imitate the actin polymerization in the arms of the phagocytic cup^31^. Similar to F-actin waves in neutrophils^40^, these structures propagate along the ventral plasma membrane, with actin polymerization at their front followed by depolymerization at their rear. In macrophages performing frustrated phagocytosis, these waves are composed of numerous F-actin puncta and undergo spatial oscillations^31^. Once the macrophage is spread, this wave may travel inward causing the cell edges to retract or move outward extending the cell boundary by forming a lamellipodium. Using RAW macrophages transfected with fluorescent myo1e/f, we found that both proteins colocalize (Supplementary Fig. 4a) with the punctate actin that makes up the wave (Fig. 3a-b, Supplementary Movie 3). We observed the same localization using primary macrophages, probing for endogenous myo1e (Fig. 3d). While myo1e/f appear to colocalize with actin punctae in the wave in xy confocal sections, xz projections reveal myo1e/f to be concentrated at the base of these structures (Fig. 3b-c). We tested whether these myosin 1-actin puncta were associated with FcR clusters by transfecting cells with EGFP-FcgRIIA. As has been previously observed, macrophages spreading on IgG formed discrete FcR clusters^41, 42^. We found that myo1f and FcgRIIA distinctly colocalized at these structures (Fig. 3e, Supplementary Movie 4), suggesting myo1e/f play a specific role at the membrane at FcR-IgG adhesions.

**Fig. 3:**
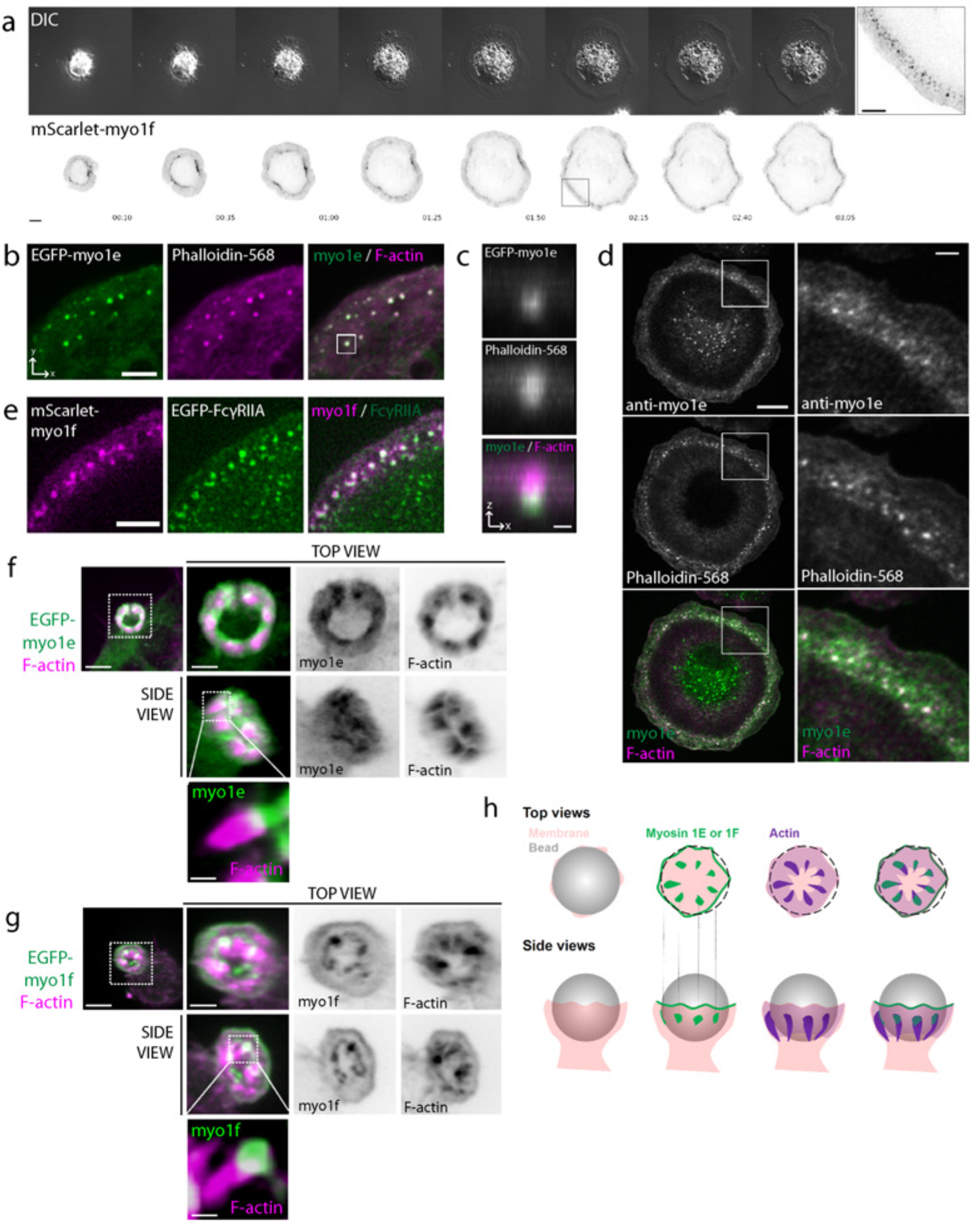
Myole and myolf localize to FcR-actin adhesions during frustrated phagocytosis. a) Time-lapse montage of RAW macrophage transfected with mScarlet-myo1f conducting frustrated phagocytosis and imaged by DIC and TIRF microscopy. Scale bar, 10 μm. Zoom of boxed region depicting myo1f puncta is shown on the right. Scale bar, 5 μm. See also Supplementary Movie 3. b) F-actin colocalizes with myo1e at F-actin puncta. Confocal section of spreading edge of RAW macrophage expressing EGFP-myo1e and stained with Alexa-Fluor-568-phalloidin. Scale bar, 5 μm. See also Supplementary Fig. 4a. c) xz projection of the boxed region in (b) showing myo1e at the base of the phagocytic adhesion. Scale bar, 0.5 μm. d) Primary macrophages form myo1e-enriched actin punctae during frustrated phagocytosis. WT BMDM were allowed to spread on IgG-coated coverslips for 10 minutes, fixed, and stained with myo1e antibody and fluorescently-labeled phalloidin. Scale bar, 10 μm. Zoom of boxed region shown on the right. Scale bar, 2 μm.e) Myo1f puncta colocalize with Fc receptors. TIRF image of the spreading edge of RAW macrophage co-expressing mScarlet-myo1f and EGFP-FcγRIIA. Scale bar, 5 μm. See also Supplementary Movie 4. f & g) Phagocytic cups contain plumes of F-actin emanating from discrete myosin 1 adhesion sites. 3D representations of phagocytic cup of RAW macrophages transfected with either EGFP-myo1e (f) or EGFP-myo1f (g) and counterstained with fluorescently labeled phalloidin (bead not outlined). Scale bar, 5 μm; inset scale bar, 2 μm; zoom scale bar, 250 nm. See also Supplementary Movie 6. h) Graphical representation of myo1e/f and F-actin localization at the phagocytic cup (top view and side view). For the top view, gray bead has been replaced by dotted outline to allow visualization inside the cup.

This is the first report of myosin-Is as components of FcR adhesions. Macrophages express multiple myosin 1 isoforms, which have also been implicated in phagocytosis (myo1g) or detected on macrophage phagosomes (myo1c)^43, 44^. Myo1c and myo1g are known as short-tailed myosins as they lack the additional tail domains of myo1e/f (TH2 and SH3 domains, see Supplementary Fig. 4f). We thus set out to test whether the localization of myo1e/f at FcR adhesions was unique compared to short-tailed myosins or reflective of a common role for myosin-Is in phagocytosis. In transfected RAW cells, neither myo1c nor myo1g localized to FcR adhesions during frustrated phagocytosis (Supplementary Fig. 4b&d), and their localization at the phagocytic cup differed from that of the long-tailed myosins, myo1e/f (Supplementary Fig. 4c&e). To test whether the additional tail domains of myo1e/f (TH2 and SH3) were responsible for their specific localization to the actin wave, we used a series of truncated myosin constructs and found that myo1e/f localization was dependent on the presence of the TH2 domain in the tail (Supplementary Fig. 4g-h). In addition, a point mutation that disrupts motor domain function also prevented myosin localization (Supplementary Fig. 4g-h;^45^). Thus, long-tailed myosins (myo1e/f) appear to have a distinctive role at FcR adhesions.

Although Fc receptors are not classically defined as adhesion molecules, the actin wave observed during frustrated phagocytosis or the inside of the phagocytic cup are undoubtedly adhesive structures. IgG coating promotes phagocyte adhesion to surfaces^46, 47^ and the tight seal created at the leading edge of macrophages performing frustrated phagocytosis is believed to create a closed compartment for cytolytic activity^48^. To test whether the actin wave of macrophages performing frustrated phagocytosis is adhesive, we attempted to detach cells exhibiting actin waves using a micropipette (see Supplementary Movie 5). Using RAW macrophages transfected with Lifeact to label F-actin, we found that cells exhibiting actin waves were considerably harder to detach and appeared attached to the substrate by the adhesions located within the wave. By comparison, transfected cells that were not forming an actin wave were easily detached. This observation supports the adhesive nature of the waves consisting of FcR-myo1e/f-actin punctae. In addition to detecting myo1e/f concentrated in punctate adhesions on planar IgG-coated surfaces, we also observed myo1e/f in distinct punctate structures at the bead-membrane interface within 3D phagocytic cups (Fig. 3f-g, Supplementary Movie 6). Extending from these myosin-I puncta were plumes of actin polymerization. We hypothesize that these myo1e/f-occupied regions represent distinct FcR adhesion sites within the cup, from which actin polymerizes to support cup structure (Fig. 3h).

### Macrophages lacking myole and myolf form disorganized actin-based adhesions during frustrated phagocytosis and exhibit denser phagocytic cups

To determine the role of myo1e/f at FcR adhesions, we conducted the frustrated phagocytosis assay using primary macrophages lacking myo1e/f. Similar to the RAW cells, BMDM spread on IgG-coated coverslips and formed circular actin waves cleared of prominent central actin. Strikingly, however, we found that actin waves in the dKO macrophages appeared thicker and more clumped (Fig. 4a).

**Fig. 4:**
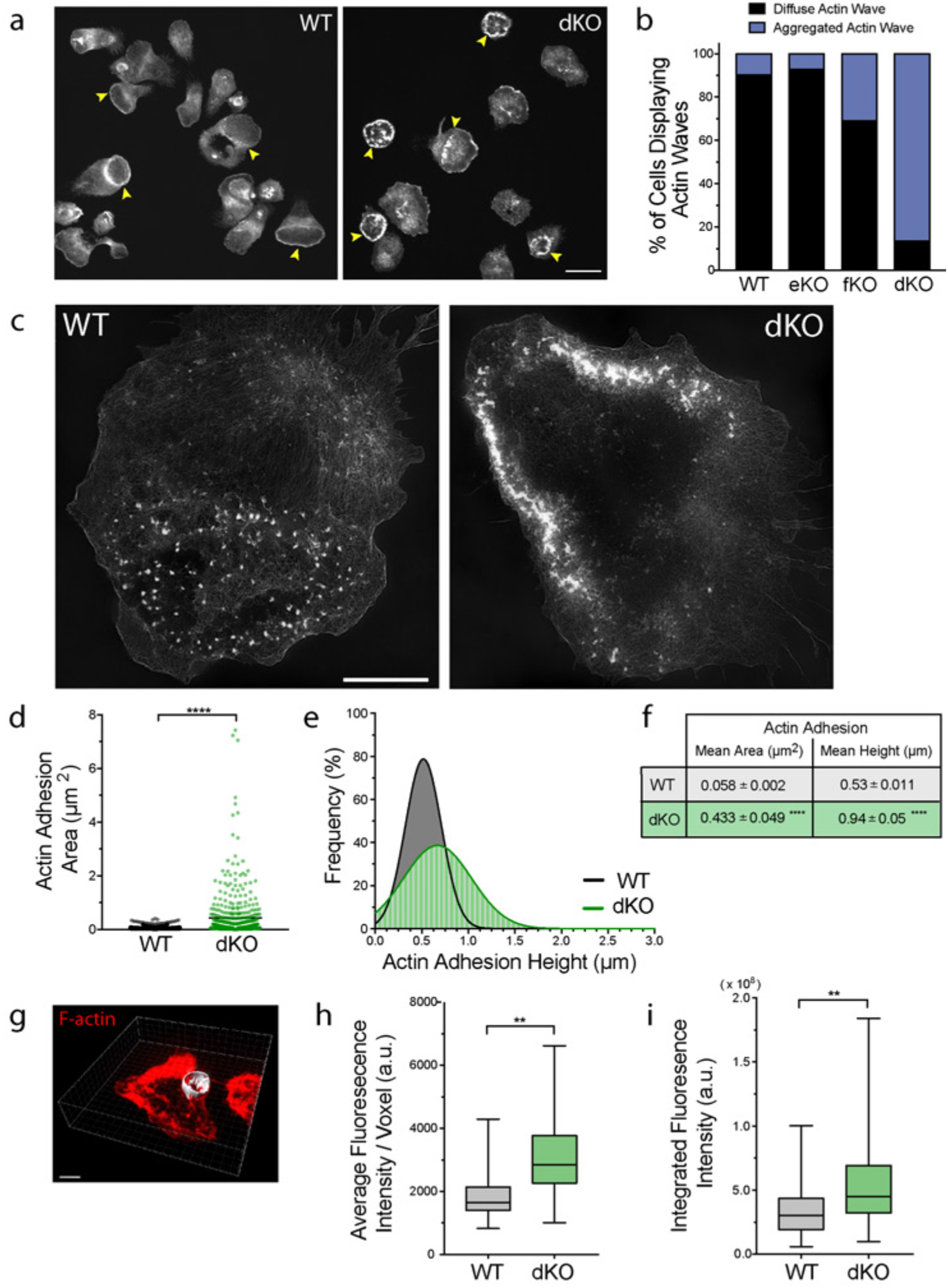
Macrophages lacking myole and myolf form disorganized actin adhesions during frustrated phagocytosis and exhibit denser phagocytic cups. a) Representative confocal image of WT and dKO macrophages performing frustrated phagocytosis. WT and dKO BMDM were allowed to spread on IgG-coated coverslips for 10 minutes. Cells were then fixed and stained with fluorescently-labeled phalloidin. Yellow arrows point to circular actin waves. Scale bar, 20 μm. b) Graph depicting the mean percentage of cells forming diffuse or aggregated actin waves in WT, myo1e^-/-^ (eKO), myof (fKO), and myo1e^-/-^; myo1f^-/-^ (dKO) BMDM. Data from 4 independent experiments (n = > 100 cells, judged blindly). See also Supplementary Fig. 5. c) Representative structured illumination microscopy (SIM) image of actin wave in WT and dKO macrophage. Scale bar, 15 μm. d) Area of individual actin adhesions in WT and dKO macrophages measured using 3D-SIM. Data from 2 independent experiments (n = > 500 adhesions from at least 14 cells per genotype, ****= p<0.0001). e) Frequency distribution of actin adhesion height in WT and dKO macrophages measured by 3D-SIM. Data from 2 independent experiments (n = > 200 adhesions from at least 15 cells per genotype). f) Table showing mean ± SEM values of actin adhesion area and height from graphs (d) and (e) (****= p<0.0001). g) Representative image of segmented phagocytic cup for fluorescence intensity measurement of Factin. WT and dKO macrophages were challenged with 6 μm IgG-coated bead, fixed and stained with fluorescently-labeled phalloidin. Phagocytic cup was detected by 3D reconstruction using Imaris software. Scale bar, 5 μm. h & i) Mean fluorescence intensity (h) and integrated fluorescence intensity (i) of actin in phagocytic cups of WT and dKO macrophages. Box and whisker plot shows median, 25^th^ and 75^th^ percentile, with error bars depicting maximum and minimum data points. Data from 3 independent experiments (n = > 100 cups per genotype, **= p<0.01).

We observed no difference in the fraction of WT and dKO cells that formed actin waves (Supplementary Fig. 5a), yet the majority of dKO cells assembled waves of clumped or aggregated actin, which were rarely observed in WT or single KO macrophages (Fig. 4b). In order to examine the actin waves at higher resolution, we used structured illumination microscopy (SIM). SIM revealed actin waves of WT macrophages to be composed of fairly uniformly sized actin punctae, similar to the actin adhesion clusters in the RAW macrophages, and reminiscent of macrophage podosomes^49^. This delicate organization was completely lost in the absence of myo1e/f, as dKO cells formed waves of clumped and aggregated F-actin (Fig. 4c). Using 3D-SIM, we measured these structures and found that actin adhesion clusters in the dKO cells were not only significantly larger in area, but also in height (Fig. 4d-f). To verify that this aggregated actin phenotype was not specific to the frustrated phagocytosis assay, we quantified actin fluorescence in fixed phagocytic cups of WT and dKO macrophages that were challenged to engulf 6 μm IgG-coated latex beads. Similar to the frustrated phagocytosis data, macrophages lacking myo1e/f assembled phagocytic cups with simply more actin (Fig. 4g-i). In an effort to rescue the aggregated actin wave phenotype of the dKO macrophages, we treated cells with small doses of Latrunculin A, which prevents F-actin polymerization by sequestering G-actin. However, such efforts proved unsuccessful, with increasing drug concentration leading only to the inhibition of cell spreading (Supplementary Fig. 5b-c). We also hypothesized that the use of Jasplakinolide might stabilize the actin waves of WT macrophages to phenocopy the actin wave morphology in the dKO cells, yet this was also not observed (Supplementary Fig. 5b-c).

It is generally accepted that the F-actin within the phagocytic cup is primarily nucleated by the Arp2/3 complex, although evidence of formin-assisted nucleation also exists^50–52^. Indeed, staining the actin waves with Arp3 antibody showed discrete puncta in WT macrophages and significant Arp2/3 recruitment to the actin clumps of the dKO macrophages (Fig. 5a). Treating dKO macrophages with CK666, an Arp2/3 inhibitor, produced a partial rescue of actin wave morphology (Fig. 5b). Given the abnormal appearance of the phagocytic F-actin in the absence of myo1e/f, we were interested in examining its supramolecular organization. We therefore performed correlative platinum replica electron microscopy (PREM) of WT and dKO macrophages undergoing frustrated phagocytosis. In control cells, the actin wave observed by PREM appeared as small clusters of branched and unbranched actin (Fig. 5c-g, Supplementary Fig. 6a and Supplementary Movie 7). In the case of dKO cells, PREM revealed a much greater density of branched and unbranched actin, elevated to a significant height above the central surface (Fig. 5h-l, Supplementary Fig. 6b and Supplementary Movie 7). We observed no apparent difference in the relative abundance of branched to linear actin within the actin waves of the WT and dKO cells. Taken together, these results implicate myo1e/f as negative regulators of Arp2/3-mediated actin polymerization within the phagocytic cup.

**Fig. 5:**
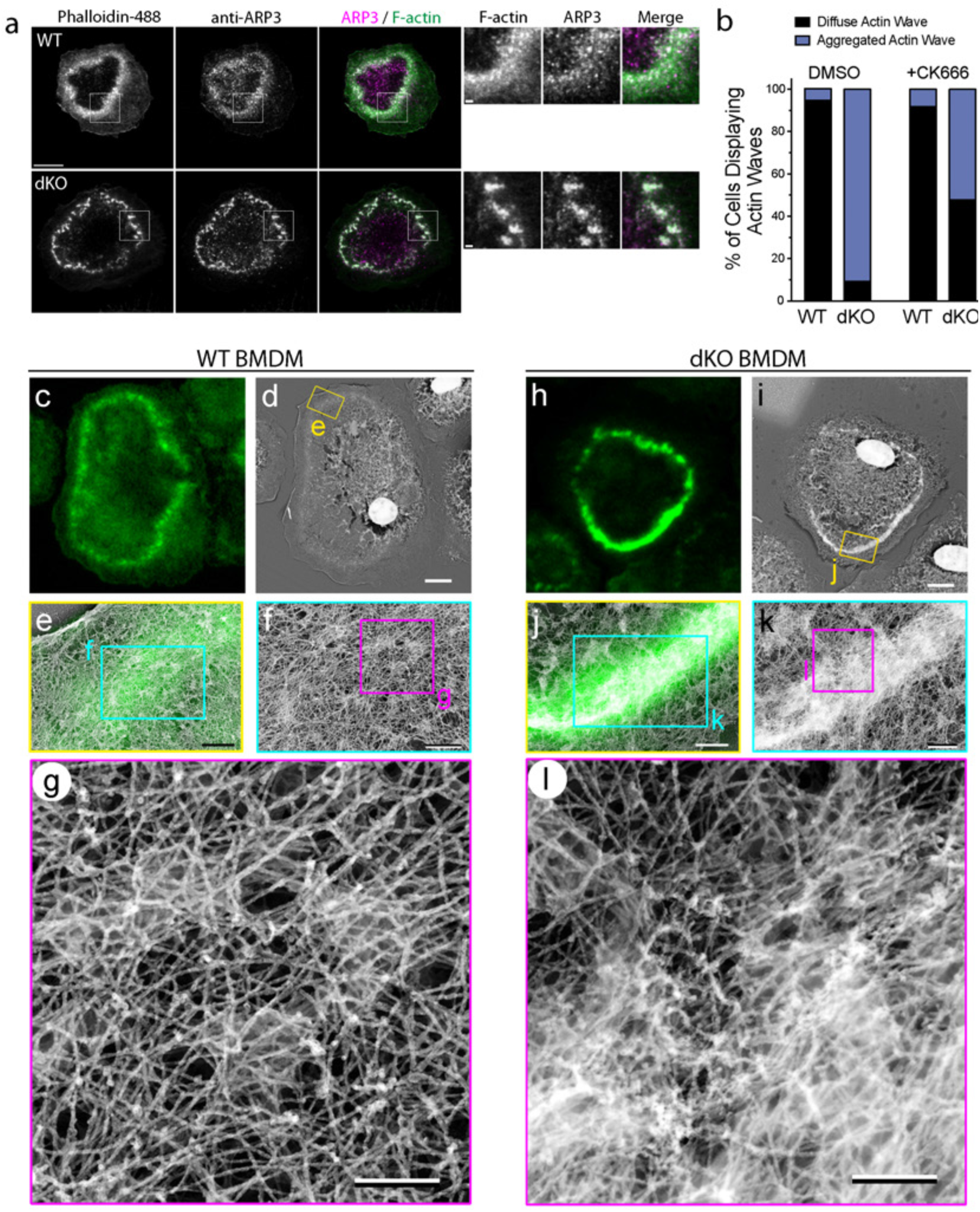
Correlative fluorescence and platinum replica EM reveals higher density of branched actin filaments in actin waves of dKO macrophages. a) Immunostaining of Arp2/3 complex in WT and dKO BMDM during frustrated phagocytosis. Cells were stained with fluorescently-labeled phalloidin and Arp3 antibody, zoom of boxed regions shown on the right. Scale bar, 10 μm; zoom panel scale bar, 1 μm. b) Treating dKO cells with low doses of Arp2/3 inhibitor CK-666 partially rescues actin wave morphology. Graph depicting the mean percentage of cells forming diffuse or aggregated actin waves of WT and dKO BMDM in the presence of 1 μm CK-666. Data from 2 independent experiments (n= >40 cells per genotype, judged blindly). c -g) Correlative confocal and platinum replica EM of actin wave in a representative WT macrophage. c) Confocal section of a WT cell stained with fluorescently-labeled phalloidin for correlative fluorescence image. Scale bar, 5 μm. d) Platinum replica EM image of the macrophage shown in (c). Scale bar, 5 μm. e) Overlay of the enlarged confocal (green) and platinum replica EM (gray) images corresponding to the boxed region in (d). Scale bar, 0.5 μm. (f & g) Sequential magnifications of boxed regions in (e) and (f), respectively, showing supramolecular architecture of individual phagocytic adhesions. Scale bars, 0.2 μm. h - l) Correlative confocal and platinum replica EM of actin wave in a representative dKO macrophage. h) Confocal section of a dKO cell stained with fluorescently-labeled phalloidin for correlative fluorescence image. Scale bars, 5 μm. i) Platinum replica EM image of the macrophage shown in (h). Scale bars, 5 μm. j) Overlay of the enlarged confocal (green) and platinum replica EM (gray) images corresponding to the boxed region in (i). Scale bars, 0.5 μm. k & l) Sequential magnifications of boxed regions in (j) and (k), respectively, showing unbridled actin polymerization. In (l), image contrast is changed by adjusting gamma. Scale bars, 0.2 μm.

### Myo1e and myo1f regulate actin dynamics at phagocytic adhesion sites, restricting adhesion size and promoting faster disassembly

Using live-cell imaging of frustrated phagocytosis by TIRFM, we were able to observe distinctly different actin dynamics between WT and dKO cells. The actin wave of WT cells moved in a sweeping fashion, with new adhesions forming quickly. These adhesions remained uniformly small and were rapidly disassembled during wave movement or collapse (Fig. 6a-b, Supplementary Movie 8). Conversely, the actin adhesion sites in the dKO macrophages moved much more slowly, with distinct fusion and fission events between adhesions leading to their clumped appearance (Fig. 6a-b, Supplementary Movie 8). This was particularly clear during wave stabilization or disassembly in which the formation of giant adhesions seemed to impair wave progression (Fig. 6b). To measure actin turnover rates within the phagocytic adhesions, we used fluorescence recovery after photobleaching (FRAP) on WT and dKO macrophages expressing EGFP-actin (Fig. 6c-d). We observed that in the presence of myo1e/f, actin was more dynamic, recovering to 86% of pre-bleached values. The immobile fraction of actin in the dKO cells was almost 3 times higher than that of WT macrophages (Fig. 6f). This indicated that the clumped actin in dKO cells was largely composed of stable actin filaments.

To better understand how the actin wave dynamics in the absence of myo1e/f results in the broad distribution of adhesion sizes observed in dKO cells (Fig. 4d), we tracked the actin wave of a dKO macrophage, quantifying both local boundary speed of the wave edge and the adjacent actin fluorescence intensity (Fig. 6g). By plotting wave boundary speed as a function of actin intensity, we observed an interesting bell-shaped relationship (Fig. 6h-l). Lower actin intensities were associated with faster protrusive wave speeds. As we observed such behavior, albeit sparingly, in the dKO cell, it seemed this fast wave expansion was unaffected by the lack of myo1e/f. However, when we tracked slower wave speeds (protrusion or retraction below 100 nm per second), we observed an inverse relationship between actin intensity in the wave and boundary speed. Strikingly, the majority of our measurements (Fig. 6h) were centered on a stalled wave, which correlated with peak actin adhesion intensities. This illustrates how the lack of myo1e/f, through the formation of large, intense and non-moving adhesive patches, impairs actin wave motility. We propose that this behavior of phagocytic adhesions in dKO cells is detrimental for phagocytic efficiency as macrophages will be unable to disassemble adhesions to finish engulfment due to overgrown adhesions that stay “glued” to the target. This would explain the observed delay for bead engulfment (Fig. 1h) as well as the actin-dense phagocytic cups of dKO macrophages (Fig. 4h).

**Fig. 6:**
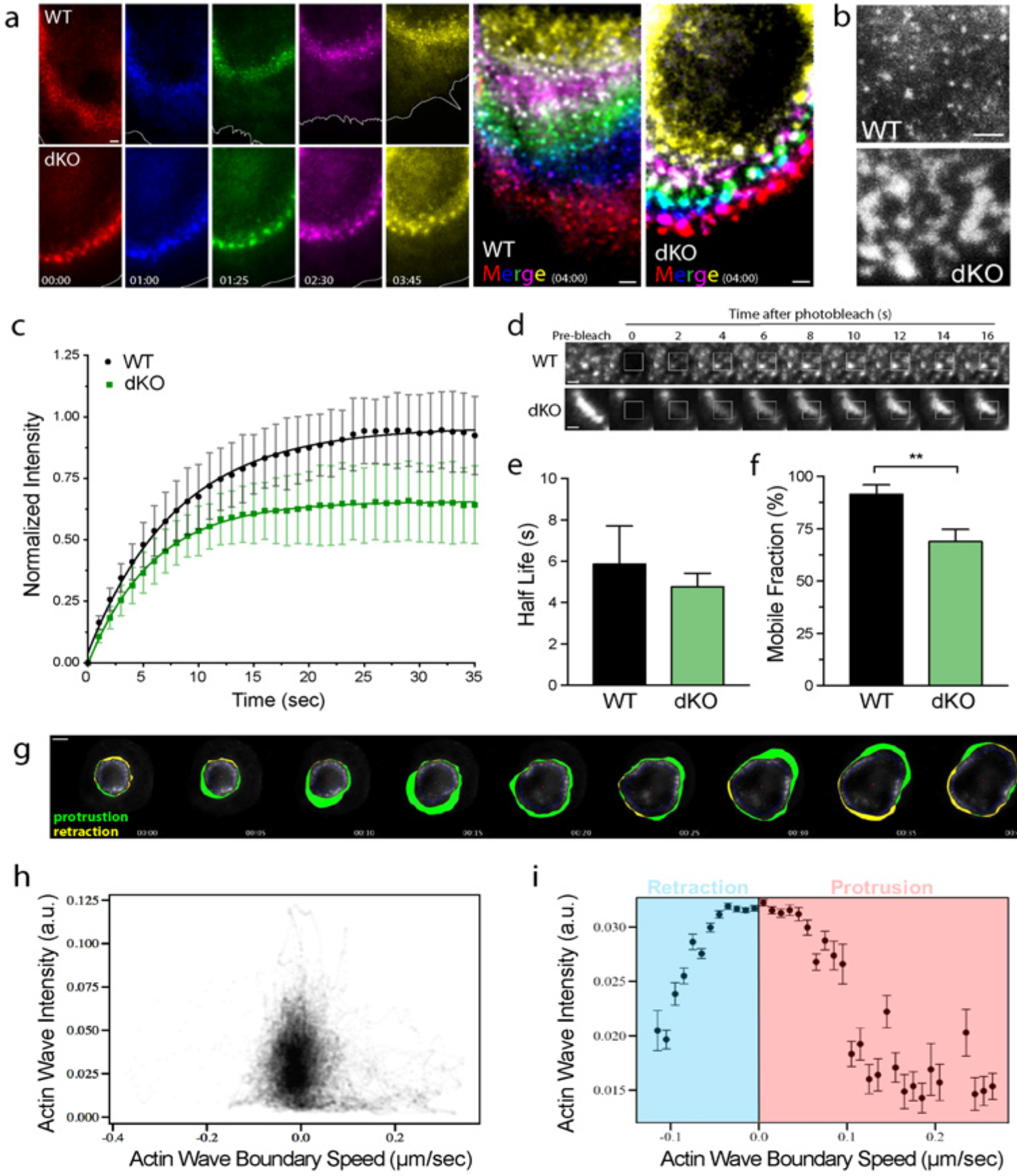
Myo1e and myo1f regulate actin dynamics at phagocytic adhesion sites, restricting adhesion size and promoting faster disassembly. a) Dynamics of actin waves in WT and dKO macrophages performing frustrated phagocytosis. BMDM were transfected with EGFP-actin and imaged by TIRFM. Montage has been color-coded with respect to time to display adhesion turnover. The white line marks the cell’s edge. Elapsed time for both time lapse images is shown at the lower left in minutes:seconds. The final images at the right (“Merge”) are maximum intensity projections of the color-coded montages. Scale bars, 2 μm. See also Supplementary Movie 8. b) Gray scale zoom of maximum intensity projections of montages in (a) to show distinct punctate structure of WT adhesions compared to fused/enlarged adhesions in dKO macrophage. Scale bar, 2 μm. c) WT and dKO macrophages transfected with EGFP-actin were subjected to FRAP analysis to measure actin turnover during frustrated phagocytosis. The resulting data was corrected for photobleaching, normalized to the pre-bleached images, then fit using the exponential functions: y = y0 + a • (1 – e^-bt^). Data from 3 independent experiments (n = 15–20 cells per genotype). d) Representative actin wave recovery as assessed by FRAP. White box indicates bleached region. Scale bar, 1 μm. e) Graph of half-life of recovery (mean ± SD). Half-life of recovery for each curve was calculated using the formula t1/2 = ln0.5/-b, where b was obtained from the exponential curve fit (p = 0.38). f) Graph of mobile fraction (mean ± SD). Percentage of recovery (mobile fraction) was calculated using formula: Xm = F_∞_ / Fi. Where Fi denotes the average fluorescence intensity before photobleaching for each normalized curve, F_∞_ refers to the average fluorescence intensity derived from the plateau for each normalized curve (**= p<0.01). g) Time-lapse montage of tracking analysis used to analyze dKO macrophage actin dynamics. dKO macrophage expressing EGFP-actin was imaged by TIRFM. The border of the actin wave was tracked for protrusion (green) / retraction (yellow) speed and the inner blue line marks the inner boundary for actin quantification. Time stamp at lower right in minutes:seconds. Scale bar, 5 μm. h) Cloud graph showing the distribution of intensities of actin adhesions vs. actin wave boundary speed during frustrated phagocytosis in dKO BMDM. dKO BMDM were transfected with EGFP-actin and imaged by TIRFM. i) Averages and standard deviation of data points depicted in (h) showing intensity of actin wave with respect to protrusion or retraction speed in dKO macrophage.

### Myo1e and myo1f control membrane mechanical properties, promoting local membrane lifting around phagocytic adhesions sites to deter adhesion sliding and expansion

To address how myo1e/f restrict adhesion size expansion to promote faster turnover and efficient phagocytosis, we focused our attention on the core function of myosin-I: their mechanical role at the interface between membrane and the actin cytoskeleton.

As membrane-actin linker, myosin-I can potentially regulate cortical tension (a mostly cytoskeleton-dependent property) as well as membrane tension, which is regulated in part by proteins that link the plasma membrane to the underlying actin cortex^53, 54^. To test this, we used atomic force microscopy (AFM) to probe the cell stiffness of WT and dKO macrophages. These AFM measurements revealed dKO macrophages to be significantly softer (Fig. 7a). As cell stiffness encompasses both cortical tension and membrane tension, we set out to evaluate the contribution of myo1e/f to membrane tension with the tether-pulling assay using optical tweezers^55, 56^. The tether force of dKO macrophages was significantly lower than that of WT macrophages, identifying myo1e/f as contributors to membrane tension in macrophages (Fig. 7b). We previously reported that macrophages undergoing phagocytosis experienced ∼30% increase in membrane tension^9^. Therefore, we decided to measure membrane tension while cells were undergoing phagocytosis. Using the optical trap, IgG-opsonized beads were placed in contact with cells to initiate phagocytosis, followed by a laser trap tether force measurement using another smaller bead coated with concanavalin A (Fig. 7c). Using primary WT BMDM, we also observed an increase in membrane tension during phagocytosis compared to resting cells. However, no such increase occurred in dKO macrophages (Fig. 7b), suggesting myo1e/f actively control membrane tension over the course of internalization.

During phagocytosis, we observed myo1e/f at two main locations: the leading edge of the phagocytic cup and phagocytic adhesion sites (Fig. 3h). The force of actin polymerization during leading edge protrusion has been described as a key contributor to membrane tension regulation in three different cell systems including macrophages^9, 57, 58^. However, we detected no difference in leading edge velocity during frustrated phagocytosis in WT and dKO macrophages. We next questioned whether myo1e/f might be regulating membrane tension locally at FcR adhesion sites, and, at the same time, restricting adhesion expansion and lateral sliding.

While imaging actin dynamics during frustrated phagocytosis by TIRFM, we noticed an interesting characteristic of phagocytic adhesions. In cells where the plasma membrane was fluorescently labeled, FcR adhesion sites were often surrounded by a circular area devoid of the membrane fluorescent signal, suggesting that the ventral surface of the cell in this circular area was located above the TIRF plane of excitation (Fig. 7d). Imaging the membrane by both TIRFM and epifluorescence proved this lifting to be specific to the ventral cell surface. This phenomenon was also observed in primary BMDM (Fig. 7e), and we hypothesized that it could be the result of a myo1e/f dependent membrane lifting away from the substrate around sites of FcR adhesions. In agreement with this hypothesis, no membrane lifting was observed at the aggregated actin adhesions in the dKO macrophages (Fig. 7e). Systematic line scans of individual adhesions among numerous cells showed this finding to be consistent (Fig. 7f-g).

**Fig. 7:**
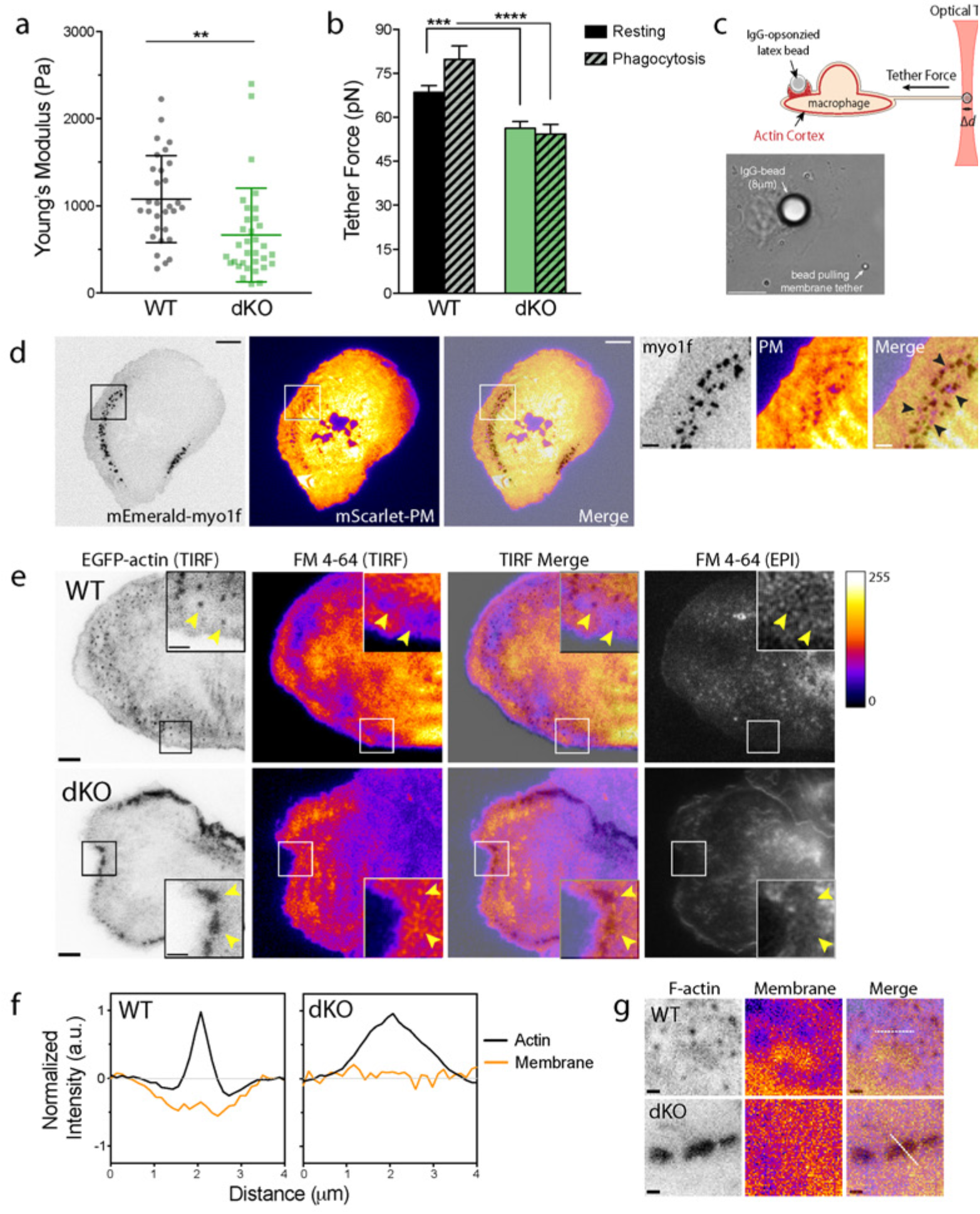
Myo1e and myo1f control membrane mechanical properties, promoting local membrane lifting around phagocytic adhesions sites. a) Myo1e/f contribute to cytoskeletal stiffness. Graph of cytoskeletal tension of WT and dKO macrophages (mean ± SD) measured using atomic force microscopy (AFM) (>30 cells measured per genotype, **= p<0.01). b) Myo1e/f promote membrane tension during phagocytosis. Membrane tension (mean ± SEM) was measured in WT and dKO macrophages while “resting” (>65 cells per genotype from 2 independent experiments, ***= p<0.001) and performing phagocytosis (>45 cells per genotype from 2 independent experiments, ****= p<0.0001). c) Schematic of experimental set-up to measure membrane tension by the tether-pulling assay during phagocytosis – corresponds to data shown in (b). Beneath, representative bright field image of tether pulling assay during phagocytosis. Scale bar, 10 μm. d) Membrane lifting observed around sites of phagocytic adhesions. RAW macrophage co-transfected with mEmerald-myo1f and mScarlet-PM (plasma membrane marker) performing frustrated phagocytosis and imaged by TIRFM. Myo1f channel has been inverted and PM channel is represented using Fire LUT. Zoom of the boxed region shown on the right denotes sites of membrane lifting around phagocytic adhesions using black arrows. Scale bar, 10 μm; zoom scale bar, 2 μm. e) Membrane lifting around sites of phagocytic adhesions (denoted by yellow arrowheads) observed in WT, but not dKO primary macrophages. WT and dKO macrophages were transfected with EGFP-actin and stained with FM 4–64 as a membrane label prior to imaging. Cells performing frustrated phagocytosis were imaged by TIRF and epifluorescence. EGFP-actin channel has been inverted and PM channel is represented using Fire LUT. Scale bar, 5 μm; inset scale bar, 2 μm. f) Average line scans of phagocytic adhesions of WT and dKO macrophages showing membrane localization with respect to F-actin by TIRFM. 4 μm line scans were performed on > 80 phagocytic adhesions in WT and dKO macrophages, transfected with EGFP-actin and stained with FM 4–64, performing frustrated phagocytosis. F-actin and membrane fluorescence intensity values were background subtracted (area around adhesion sites) and normalized to peak intensity values. Data from 3 independent experiments (>15 cells analyzed per genotype). g) Representative example of drawn line scans (dashed white line) at phagocytic adhesions of WT and dKO macrophages performing frustrated phagocytosis. EGFP-actin channel (F-actin) has been inverted and FM 4–64 channel (membrane) is represented using Fire LUT. Corresponds to data in (f). Scale bar, 1 μm.

In light of these results and in combination with our AFM and tether measurements, we propose that myo1e/f are tethering the plasma membrane at individual adhesion sites, preventing undesirable adhesion overgrowth by locally lifting the membrane around adhesions, and thus regulating membrane tension. This adhesion restriction by membrane lifting enables productive actin polymerization and adhesion site turnover required for proper phagocytosis.

## Discussion

Here we show that both myo1e/f are required for efficient FcR-mediated phagocytosis. Compared to other proteins involved in phagocytic ingestion, myo1e/f are uniquely localized at the tip of the phagocytic cup, preceding actin polymerization. We observed myo1e/f between F-actin and membrane at FcR adhesions, with myo1e/f being located slightly more ventrally than actin, in both the 2D frustrated phagocytosis assay and inside the 3D phagocytic cup. Using deletional analysis, we discovered that localization of myo1e/f to the actin wave (or phagocytic cup) depended partly on the TH2 domain in the tail and a functional motor domain. This demonstrates a specific role for long-tailed myosinIs in phagocytosis, as short-tailed myosin-Is do not contain this tail region. These findings parallel myosin-I behavior reported in *Dictyostelium*, in which planar actin waves are also used as a model for the phagocytic cup^59,60^. The TH2 domain of long-tailed myosin-Is contains an ATP-insensitive actin-binding site^61–63^ as well as a large number of basic amino acids that could bind to membrane phospholipids^16^. Thus, it appears that myo1e/f recruitment to the phagocytic cup may depend in part on actin binding via the motor domain and in part on the interactions of the TH2 domain with actin filaments or membrane phospholipids.

The process of phagocytosis involves several steps that could potentially rely on myosin-I functions. Extension of pseudopods during initial cup formation, focal exocytosis to increase cup surface area, and contraction of the actin filaments within the cup could all be driven by myosin activity. However, we found that none of these events appear to be connected to the activity of myo1e/f. Phagocytic cups still formed in the absence of these myosins, indicating that initial actin assembly and pseudopod extension were not disrupted. Focal exocytosis markers did not colocalize with myo1e/f, and dKO macrophages did not exhibit defects in membrane dynamics suggesting myo1e/f play no direct role in membrane addition to the cup. Unexpectedly, the loss of myo1e/f led to a dramatic change in actin dynamics during frustrated phagocytosis, resulting in excessive accumulation of branched actin networks with low turnover at FcR adhesion sites. By measuring membrane tension, we confirmed that myo1e/f function to connect the plasma membrane to the underlying actin cortex in macrophages, and that an increase in membrane tension that is normally induced by phagocytosis is not observed in myo1e/f null cells. Previous electron microscopy studies show that in cross-sections of phagocytic cups, cell-target contact sites are not continuous but rather separated by regions where the plasma membrane is lifted away from the surface of the target^64,65^. In macrophages lacking myo1e/f, this membrane lifting was not observed at sites of phagocytic adhesions, indicating that the loss of membrane-actin tethering alters membrane geometry at the adhesion sites. Based on these observations, we propose a novel role for myo1e/f in organizing membrane around actin-based adhesions during phagocytosis. We hypothesize that the loss of membrane organization at FcR adhesions disrupts adhesion turnover and alters the dynamics of F-actin polymerization within the cup. This manifests overall in the generation of phagocytic cups that have denser F-actin and complete closure/internalization at a slower rate.

One possible explanation for the excessive actin polymerization phenotype in the absence of myo1e/f is that these myosins could be necessary for the activity or localization of actin depolymerizing proteins or actin severing proteins. Macrophages lacking CapG, a protein that caps barbed ends enabling F-actin turnover, display a 50% reduction in the rate of Fc-receptor mediated phagocytosis^66^. Yet using an antibody against capping protein, we could not detect any difference in the ratio of capping protein to F-actin in WT and dKO cells conducting frustrated phagocytosis (data not shown). Moreover, myo1e has been reported to bind actin regulatory proteins such as WASP or CARMIL^67,68^, and we have found no difference in the recruitment of these proteins to the phagocytic cup or in resting macrophages lacking myo1e/f (data not shown). Intriguingly, a recent paper has reported that F-actin depolymerization rates increase when actin filaments slide on myosin 1b (myo1b)^69^. However, since the kinetics of the myo1b motor domain are distinct from that of long-tailed class 1 myosins, this feature is unlikely to be generalizable to myo1e/f. We therefore conclude that the phenotype of myo1e/f loss results solely from its biophysical role at phagocytic adhesion sites.

Myo1e/f indeed promote phagosome closure, yet not through the generation of significant contractile force as was previously assumed^26^. Using traction force microscopy, we showed that loss of myo1e/f did not affect contractile behavior during frustrated phagocytosis. However, the myosin II inhibitor blebbistatin has been shown to have a dramatic effect on macrophage traction forces using the same assay^27^. A recent study suggests myosin II promotes timely squeezing at the base of the phagocytic cup to ensure ingestion^70^. Another interesting suggestion for myosin II function is the exclusion of specific cell receptors, such as CD47 or phosphatase CD45, from the phagocytic cup to allow proper downstream signaling^71, 72^. Hence, the proposed function of myo1e/f here is independent of the roles that might be played by myosin II during phagocytosis.

Overall, our findings emphasize a heretofore unappreciated component of macrophage phagosome closure: dynamic adhesions at the bead-membrane interface, which closely couple actin polymerization and increased membrane tension to target engulfment and cup closure. While myo1e/f in macrophages regulate membrane tension globally, their concentrated action within the phagocytic cup appears to control the size and consequently dynamics of phagocytic adhesions. By lifting membrane around individual adhesions, myo1e/f prevents adhesion fusion and expansion, which results in slower internalization. Membrane tension has previously been shown to provide mechanosensitive feedback that limits or redirects F-actin polymerization during neutrophil migration^73,74^. Within lamellipodia in worm sperm cells, reduced membrane tension results in less organized cytoskeletal filaments and a slower extension speed^75^, and a similar mechanism may be involved during phagocytic cup closure. Importantly, our observations appear to provide a mechanistic, molecular basis for the “enveloping embrace” phenomenon that was first predicted in the biophysical and modeling studies of neutrophil phagocytosis^76^. These studies posit that phagocytes use myosin-I-based adhesions, which couple the internal actin cytoskeleton to the plasma membrane, to effectively anchor the bead to the cell allowing F-actin polymerization within the cup to push forward and finish internalization. One can envision myosin-Is directing actin polymerization via mechanosensitive regulation of membrane-limited adhesion sites in a variety of settings, including cell migration and contact assembly^77^. Thus, while our studies reveal novel roles for myosins in phagocytosis, these findings may also have a broader significance in other physiological settings.

## Materials and Methods

### Materials and correspondence

Requests for reagents and resources should be directed to and will be fulfilled by the lead contacts, Mira Krendel and Nils Gauthier.

### Mice

Myo1e^-/-^ mice^23^ and myo1f^-/-^ mice^24^ were crossed to create myo1e^-/-^; myo1f^-/-^ double knockout (dKO) mice. All mice used in this study were maintained on a C57BL/6 background. All procedures utilizing mice were performed according to animal protocols approved by the IACUC of SUNY Upstate Medical University. Both male and female mice were used; for each individual experiment utilizing BMDM, bone marrow preparations from age- and sex-matched mice were used.

### Cell culture

RAW264.7 (ATCC; male murine cells) were cultured in Dulbecco’s Modified Eagle Medium (DMEM), high glucose, containing 10% FBS and 1% antibiotic-antimycotic (Gibco) at 37°C with 5% CO_2_. All transfections of RAW cells and BMDM were accomplished by electroporation (Neon) using manufacturer’s instructions.

### Bone marrow isolation and primary macrophage culture

Following euthanasia, femurs and tibias of mice were removed and flushed with Dulbecco’s Modified Eagle Medium (DMEM) containing 10% FBS and 1% antibiotic-antimycotic (Gibco). Red blood cells were lysed using ACK buffer (0.15M NH_4_Cl). Bone marrow progenitor cells were recovered by centrifugation (250 × g, 5 minutes, 4°C), washed once with sterile PBS and plated on tissue culture dishes in a 37°C incubator with 5% CO_2_. The next day, non-adherent cells were moved to bacteriological (non-tissue-culture treated) Petri dishes and differentiated in DMEM, high glucose with 20% L929-cell conditioned media (v/v),10% FBS, and 1% antibiotic-antimycotic, with fresh medium given on day 3 or 4. Cells were confirmed to be macrophages (F4/80^+^ CD11b^+^) by flow cytometry. All experiments were done within five days post-differentiation.

### Chemicals and drugs

Latrunculin A, CK-666 and LY294002 were purchased from EMD Millipore. Jasplakinolide, Concanavalin A, and unlabeled phalloidin were purchased from Sigma. Alexa-Fluor-488 or Alexa-Fluor-568 conjugated phalloidin were purchased from Life Technologies and FM 1–43 or FM 4–64 were purchased from Invitrogen.

### Antibodies

The following antibodies were used in this study: rabbit anti-myo1e has been previously described^78^; myo1f (Santa Cruz, B-5, #376534); Arp3 (Millipore, clone13C9, #MABT95); BSA (Sigma, clone 3H6, #SAB5300158); rat anti-mouse CD16/32 (BD, #553141); AffiniPure mouse anti-rat (Jackson Labs, 212–005-082); the following antibodies were purchased from Cell Signaling Technologies: pSyk (#2701), Syk (#13198), pAkt (#4060), Akt (#4691), pERK (#4370), ERK (#9102); fluorescent secondary antibodies against mouse or rabbit (Life Technologies).

### Constructs

Human myosin 1e and truncated constructs tagged with GFP have been previously described^79^. Human myosin 1f (NM_012335.3) cDNA from the Mammalian Gene Consortium was amplified and cloned into pEGFPC1 (Clontech). Both myosins were also cloned into mEmerald-C1 and tdTomato-C1 (gifts from Michael Davidson; Addgene plasmids #53975 and 54653) and mScarlet-C1. The following constructs were purchased from Addgene: EGFP-PLCδ-PH, EGFP-AKT-PH and EGFP-PKCd-C1 (gifts from Tobias Meyer, plasmids #211789, 21218, and 21216), mScarlet-PM (gift from Dorus Gadella, #98821), EGFP-VAMP3 (a gift from Thierry Galli, #42310), Ruby-Lifeact, mEGFP-Lifeact, and EGFP-actin (gifts from Michael Davidson, #54674, #54610, and #56421). EGFP-FcgRIIA was a gift from Sergio Grinstein (Hospital for Sick Kids, Toronto, ON). YFP-myo1c and EGFP-myo1g were gifts from Matt Tyska (Vanderbilt University, Nashville, TN) that were cloned into mEmerald-C1.

### Immunostaining

Cells were fixed using fresh 4% paraformaldehyde/PBS for 15 minutes. After washing away fixative, cells were permeabilized in 0.1% Triton X-100/PBS for 3 minutes. Cells were blocked for 30 minutes at room temperature with 5% normal goat serum/ 3% BSA dissolved in PBS with 0.05% Tween-20. Cells were exposed to primary antibodies at the appropriate dilutions for one hour at room temperature. Cells were then washed 3 times for 5 minutes with PBS/0.05% Tween-20. Secondary antibodies and fluorescent phalloidin were then applied for 30 minutes at room temperature. Cells were then washed again for 3 times, 5 minutes each before mounting with Prolong Diamond Antifade Mountant (Invitrogen).

### FcR stimulation

Fc receptor stimulation was conducted as previously described^80,81^ with a few alterations. BMDM were plated at 6 × 10^6^ cells per 10 cm petri dish. Cells were serum-starved for 6 hours and then incubated with 10 μg/mL rat anti-mouse CD16/32 antibody (2.4G) in cold serum-free media for 40 minutes at 4°C to allow binding. Cells were then quickly washed 2X with ice-cold PBS to remove unbound antibody and exposed to warm serum-free media containing 15 μg/mL anti-rat antibody to initiate crosslinking. At the specified time points, crosslinking media was removed and the cells were processed for western blot analysis. The “0” time point control dish was left at 4°C.

### Western blotting

Cells were washed 1X in ice-cold PBS and harvested by scraping in NP-40 lysis buffer (1% NP-40, 150 mM NaCl, 50 mM Tris-HCl, 10 mM NaF) with phosphatase and protease inhibitors (Roche). Cells were rotated at 4°C for 25 minutes and then pelleted at 16,000 × g for 15 minutes at 4°C. Supernatant was then removed and boiled with Laemmli sample buffer, and separated on 10–20% gradient SDS-PAGE gel, followed by transfer to PDVF. Membranes were blocked in 5% milk or 3% BSA in TBST for one hour at room temperature. Primary antibodies were diluted in 5% milk or 3% BSA in TBST and incubated with the membrane overnight at 4°C. The next day, the membrane was washed 3X for 5 minutes in TBST. HRP-conjugated secondary antibodies were diluted in 5% milk or 3% BSA in TBST and exposed to membranes for one hour at room temperature. Chemiluminescence was detected using WesternBright Quantum (Advansta) and imaged on a Bio-Rad ChemiDoc imaging system.

### Phagocytosis assay

Polystyrene beads (PolySciences Inc., 2, 6, or 8 μm) were washed in PBS and opsonized overnight at 4°C in 3 mg/ml mouse IgG (Invitrogen). To remove excess antibody, beads were washed three times with PBS and resuspended in sterile PBS. Beads were applied to macrophages in a 12-well plate at an estimated ratio of 10:1. To synchronize phagocytosis, the plate was spun at 300 x g for 2 minutes at 4°C. Cells were incubated at 37°C to initiate phagocytosis. To stop phagocytosis, cells were washed three times with ice-cold PBS to remove unbound beads and fixed with 4% PFA/PBS for 15 minutes. Cells were then washed and stained with goat anti-mouse-Alexa Fluor-568 antibodies for visualization of un-internalized beads for 30 minutes. Cells were then washed with PBS (3 × 5 minutes) and permeabilized with 0.025% Triton X-100/PBS, then stained with Alexa Fluor 488-conjugated phalloidin, followed lastly by DAPI (NucBlue Fixed Cell ReadyProbe Reagent, Invitrogen). Coverslips were then mounted using Prolong Diamond Antifade Mountant. For phagocytic internalization/association quantification, 20 fields of view per sample were imaged by spinning disk confocal microscopy using 20X magnification. Internalized beads were visualized using the brightfield channel. Quantification resulted in at least 200 cells being analyzed in three independent experiments.

### Frustrated phagocytosis assay

Frustrated phagocytosis assays were conducted as previously described^31^. In brief, glass coverslips were acid-washed with 20% nitric acid. They were then coated in 1 mg/mL BSA/PBS for 1 hour at 37°C, followed by incubation with 10 μg/mL mouse anti-BSA antibody (Sigma) for 1 hour at 37°C. Coverslips were washed 3X in PBS before use. For live-cell imaging, coverslips were affixed to a custom imaging chamber with vacuum grease (Dow Corning). Prior to the assay, cells were serum starved in 1X Ringer’s buffer (150 mM NaCl, 5 mM KCl, 1 mM CaCl_2_, 1 mM MgCl_2_, 20 mM HEPES and 2 g/l glucose, pH 7.4) for 20 minutes. Cells were introduced to the mounted chamber and imaged spreading in 1X Ringer’s buffer at 37°C. In the case of drug studies, cells were exposed to the drug while in suspension and carried out frustrated phagocytosis in the same dilution of the drug.

### Flow cytometry

Macrophages were pelleted (250 × g, 5 min, 4°C, 1 × 10^6^ cells/tube) and resuspended in fresh FACS buffer (5% FBS/0.1% NaN_3_/PBS) and blocked with rat anti-mouse CD16/32 (Biolegend, clone 93, #101301) for 30 minutes on ice. Cells were then incubated with FITC-F4/80 (Biolegend, clone BM8, #123107) and APC-CD11b (BD Biosciences, clone M1/70, #553312) for 50 minutes in the dark. Cells were washed twice and resuspended in 0.2 mL FACS buffer for immediate analysis with a BD LSRII flow cytometer and BD FACSDiva program. Fc receptors were quantified using APC-CD16/32 (BD, clone 2.4G2, #558636). Data was processed using FlowJo software.

### Platinum replica electron microscopy

For correlative light and platinum replica electron microscopy, cells were allowed to spread on IgG-coated glass pre-coverslips coated with a thin layer of gold through a locator grid (400 mesh, Ted Pella, Inc., Redding, CA). The gold layer provided the coverslips with a finder grating recognizable by both light and electron microscopy^82^. The cells were extracted for 3 minutes with extraction solution containing 1% Triton X-100, 2% PEG (MW 35,000), 4 μM unlabeled phalloidin in M buffer (50 mM imidazole, pH 6.8, 50mM KCl, 0.5 mM MgCl_2_, 1mM EGTA, and 0.1 mM EDTA). Samples were triple washed in M buffer with 4 μM unlabeled phalloidin and fixed in 2% glutaraldehyde in PBS buffer for 20 minutes. Then samples were triple washed in PBS and quenched with 5 mg/mL NaBH_4_ in PBS for 10 minutes and again triple washed in PBS. Next, cells were stained with Alexa Fluor 488-phalloidin and imaged by spinning disk confocal microscopy. Samples for correlative platinum replica electron microscopy were processed as described previously^83,84^. Samples were analyzed using JEM 1011 transmission electron microscope (JEOL USA, Peabody, MA) operated at 100 kV. Images were captured by ORIUS 832.10W CCD camera (Gatan, Warrendale, PA) and presented in inverted contrast.

### Traction force microscopy (TFM)

Linearly elastic polyacrylamide gels with a shear modulus of 1.5 kPa were prepared using previously published protocols^85^, with a final mixture of 7.5% acrylamide/0.05% bis-acrylamide solutions (Bio-Rad). 40 nm dark-red fluorescent beads (Invitrogen) were included in polymerization mixture at 1:100 dilution. 6 μL of the PAA solution was pipetted onto a hydrophobic glass slide, covered with a silanized coverslip (No. 1.5, Electron Microscopy Sciences) and allowed to polymerize for 30 minutes. The gels were removed and washed in PBS. BSA (Fisher Bioreagents) was coupled to the surface of the gel using the photoactivatable crosslinker sulfo-SANPAH (Thermo Scientific). Gels were incubated in a 2 mg/mL solution of sulfo-SANPAH and exposed for 5 minutes at 5W in a UV-crosslinker (Analytik-Jena), rinsed with water, and incubated inverted on parafilm in a 1 mg/mL solution of BSA for 1 hour at room temperature. The gels were washed several times in PBS, and further incubated with a mouse anti-BSA antibody (Sigma) at 10 μg/mL for 1 hour, and finally washed thoroughly with PBS. Cells were imaged on a Nikon Ti-E Microscope with Andor DragonFly Spinning Disk Confocal system using a 60X (1.49 N.A.) TIRF Objective. Images were recorded on a Zyla 4.2 sCMOS camera. Cells were maintained at 37°C and 5% CO2 in an environmental chamber (Oko Labs). All hardware was controlled using Andor’s iQ software. Approximately 50,000 cells were added to the gels immediately prior to imaging. Traction forces were calculated as previously described^86^. Briefly, bead images in a time series were first registered using a region devoid of cells. Bead displacement in the substrate was determined using PIV software (OpenPIV; MATLAB), which compared each bead image to a reference image before the cell had attached to the substrate. Each window was 3.46 μm x 3.46 μm in size with a center to center distance of 1.73 μm. Displacement vectors were filtered and interpolated using the Kriging interpolation method. Traction stresses were reconstructed via Fourier Transform Traction Cytometry, with a regularization parameter chosen by minimizing the L2 curve. The strain energy was calculated as one half the integral of the traction stress field dotted into the displacement field^87^. Spread areas were calculated using DIC images traced by hand.

### TIRF microscopy

TIRF imaging was performed on multiple systems: (Figure 4SH) an Olympus inverted microscope equipped with an iLas2 targeted laser illuminator (Roper Scientific) using both a 63X and 100X objective (1.49 N.A.). Fluorescence was spectrally filtered and collected using a pair of Evolve EMCCD cameras (Photometrics) for red and green emission. (Figure 3E, 7D) a Nikon Eclipse TE2000-E2 multimode TIRF microscope equipped with PRIME-95B cMOS camera (Photometrics), CFI Apo 100X (1.49 N.A.) oil TIRF objective (Nikon), OKO Labs temperature-controlled microscope enclosure (OKO Labs), LUNA-4 solid state laser (Nikon). All other images were collected on: True MultiColor Laser TIRF Leica AM TIRF MC equipped with an Andor DU-885K-CSO-#VP camera and a 63X (1.47 N.A.) oil CORR TIRF objective. For actin/membrane line scans, 4 μm long line scans, centered over phagocytic adhesions, were drawn on the TIRF images of WT and dKO BMDM using ImageJ. Fluorescence intensities of F-actin and membrane markers measured along the lines were background-subtracted, with the background defined as the mean fluorescence intensity in the region directly surrounding adhesions. For comparison among cells, F-actin values were normalized to peak values (set to 1) and membrane values were normalized to the lowest value (set to –1). Average line scans were graphed with F-actin peak at center (2 μm). Over 80 adhesions were measured in over 15 cells per genotype from three independent experiments.

### Confocal microscopy

Images were taken using a PerkinElmer Ultra View VoX Spinning Disc Confocal system mounted on a Nikon Eclipse Ti-E microscope equipped with a Hamamatsu C9100–50 EMCCD camera, a 100X (1.4 N.A.) PlanApo objective, and controlled by Volocity software.

### Structured illumination microscopy

SIM images were acquired using Nikon N-SIM E microscopy system based on the Eclipse Ti research inverted microscope with CFI Apo TIRF SR 100X (1.49 N.A.) objective and Hamamatsu ORCA-Flash 4.0 V2 camera.

### Atomic force microscopy

AFM indentation was carried out using JPK NanoWizard3 mounted on an Olympus inverted microscope. The protocol was adapted from a previous study^88^. A modified AFM tip (NovaScan, USA) attached with 10 μm diameter bead was used to indent the center of the cell. The spring constant of the AFM tip cantilever is ∼0.03 N/m. AFM indentation loading rate is 0.5 Hz with a ramp size of 3 μm. AFM Indention force was set at a threshold of 2 nN. The data points below 0.5 μm indentation depth were used to calculate Young’s modulus to ensure small deformation and minimize substrate contributions^88^. The Hertz model is shown below: 
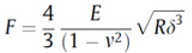
where F is the indentation force, E is the Young’s modulus to be determined, ν is the Poisson’s ratio, R is the radius of the spherical bead, and δ is indentation depth. The cell was assumed incompressible and a Poisson’s ratio of 0.5 was used.

### Phagocytic cup F-actin quantification

Phagocytic cups of WT and dKO BMDM, fixed and stained with fluorescently labeled phalloidin, were imaged using a 100X objective by spinning disk microscopy with 0.3 μm z-steps. Images were reconstructed using Imaris software and the average fluorescence intensity per voxel and integrated intensity were measured for the ROI enclosing the cup. Over 100 cups per sample were analyzed in three independent experiments.

### FM 1–43 quantification

FM 1–43 experiments were carried out as previously described^9^. Serum-starved resuspended macrophages were exposed to 10 μg/mL FM 1–43 before being added to an imaging chamber containing background 5 μg/mL FM 1–43 in 1X Ringer’s Buffer. As cells spread, DIC and FITC images were collected at 15 second intervals using a 40X (1.35 N.A.) water objective mounted on a DeltaVision microscope (Olympus IX70; Applied Precision) equipped with a CoolSNAP HQ camera (Photometrics) and SoftWoRx software. As FM 1–43 has a quantum yield 40X higher in lipid membranes, total FM 1–43 intensity from a cell at a given time point is linearly related to the sum of the initial plasma membrane area and total net membrane exocytosed. Intensities were extracted from images and background corrected using ImageJ before being normalized to the initial pre-spread intensities.

### FRAP analysis

Fluorescence recovery after photobleaching (FRAP) was performed using a Perkin Elmer UltraView VoX Spinning Disk Confocal system equipped with the Photokinesis module. Photobleaching using full power of a 488 nm argon laser was performed by selecting a square ROI. Post-bleach images were collected at 1 second intervals. Changes in ROI fluorescence intensity were measured over time using ImageJ and corrected using the background intensity and a control region of interest to account for any acquisition bleaching, and normalized to pre-bleach and post-bleach intensity values. The best fit curve for fluorescence recovery was obtained using Prism Software. The following equation was used: y = a(1-e^-bx^), where x is seconds. The half time of recovery was determined using b from the previous equation, where t_1/2_ = ln0.5/-b.

### Membrane tension measurements

Membrane tension was measured as previously described^9^. In summary, an optical tweezer was generated on a Nikon A1-R microscope by focusing a 5.0-W, 1,064-nm laser through a 100X (1.3 N.A.) objective (Nikon). 1 μm polystyrene beads (Polysciences Inc.) coated in Con A were used to pull tethers from macrophages. Bright-field images were acquired using an Andor camera. The trap strength was calibrated with the help of a previously described method^89^, by observing Brownian motion of trapped beads with an exposure time of 0.6 ms to minimize motion blur. The measured bead displacement was tracked using ImageJ and converted into measured force.

### Statistical analysis

All comparisons were carried out using an unpaired two-tailed t-test for independent samples, with differences between genotypes considered statistically significant at p < 0.05. Statistical analyses and graphing was performed by GraphPad Prism software.

### Actin wave boundary speed quantification

To analyze the actin wave boundary speed we wrote a custom macro for Fiji^90^. The macro is available upon request. Signal background was subtracted and the cell manually segmented in all frames. Once identified the cell center, for each angle between 0 and 359 degrees, the boundary speed was computed measuring the distance between the boundaries in two consecutive frames. Speed is defined positive when boundary expands and negative when it contracts. The corresponding mean actin intensity for the same angles was computed averaging the signal intensity from the boundary to 30 pixels inward (∼4um). Results were analyzed and plotted with R^91^ and the ggplot2 R package^92^.

## End Matter

### Author Contributions and Notes

S.R.B. designed and performed experiments, analyzed data, and wrote the manuscript. N.R. and P.O. performed TFM experiments and analyzed TFM data. M.S. and T.S. performed platinum replica EM studies. Q.L. assisted with AFM experimentation and analysis. P.M. helped analyze data. M.M. and R.A.F. generated mouse model. M.K. and N.G. designed experiments and wrote the manuscript. All authors reviewed the manuscript prior to publication.

The authors declare no conflict of interest.

## Acknowledgments

This work was supported by the NSF (#1515223) and AHA (18PRE34070066) grants to S.R.B., the National Institute of Diabetes and Digestive and Kidney Diseases of the NIH under Award R01DK083345 to M.K., the National Institute of General Medical Sciences grant R01GM095977 to T.S., and the Italian Association for Cancer Research (AIRC), Investigator Grant (IG) 20716 to N.G. A travel award grant from the Boehringer Ingelheim Fonds to S.R.B. also made this work possible. We would like to thank Phuson Hulamm and Nicholas Deakin, Ph.D. (Nikon Instruments Inc.) for assistance with SIM image acquisition. The authors gratefully acknowledge technical assistance from Sharon E. Chase.

---

**Supplementary Fig. 1:**
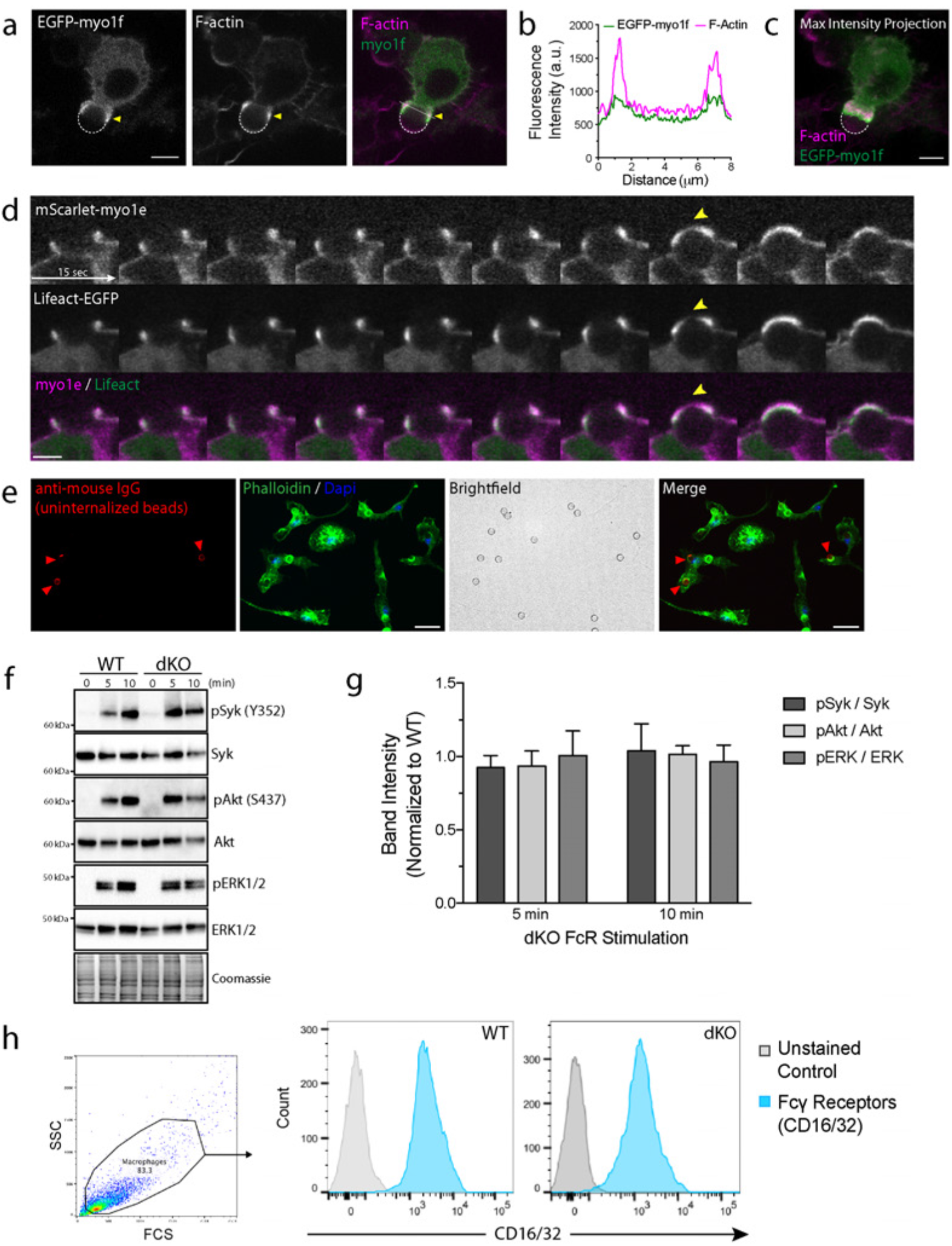
Myo1e and myo1f are required for efficient phagocytosis but do not affect phagocytic signaling. a) Representative confocal section of RAW264.7 macrophage transfected with EGFP-myo1f engulfing a 6 μm IgG-coated bead and stained with fluorescently-labeled phalloidin. Yellow arrowheads point to the phagocytic cup, and the bead is outlined by a dotted white line. Scale bar, 5 μm.. b) Line scan of EGFP-myo1f and F-actin intensities along the white line in (a) Merge to illustrate myo1f and actin colocalization at the phagocytic cup. c) Max intensity projections of (a) Merge show that myo1f precedes F-actin at the leading edge of the phagocytic cup. d) Time-lapse montage of a single confocal section obtained from RAW macrophage expressing mScarlet-myo1e and Lifeact-EGFP engulfing 8 μm IgG-coated bead. Scale bar, 5 μm. Data source is Supplementary Movie 1. e) Representative image series of phagocytosis assay to quantify internalization. Cells were stained without permeabilization using anti-mouse secondary antibody to identify un-internalized beads. This channel along with the bright field image was used to identify completely internalized beads. Cells were identified/counted using phalloidin/DAPI. Scale bar, 25 μm..f) Representative Western blot analysis of phagocytic signaling pathways induced by FcR crosslinking in WT and dKO macrophages. The 0 timepoint represents an un-induced state. g) Quantification (mean ± SEM) of Western blots used to examine FcR signaling, with band intensities for dKO macrophages normalized to WT values. Data from 3 independent experiments. h) Flow cytometry analysis of FcRs present on the cell surface in WT and dKO macrophages.

**Supplementary Fig. 2:**
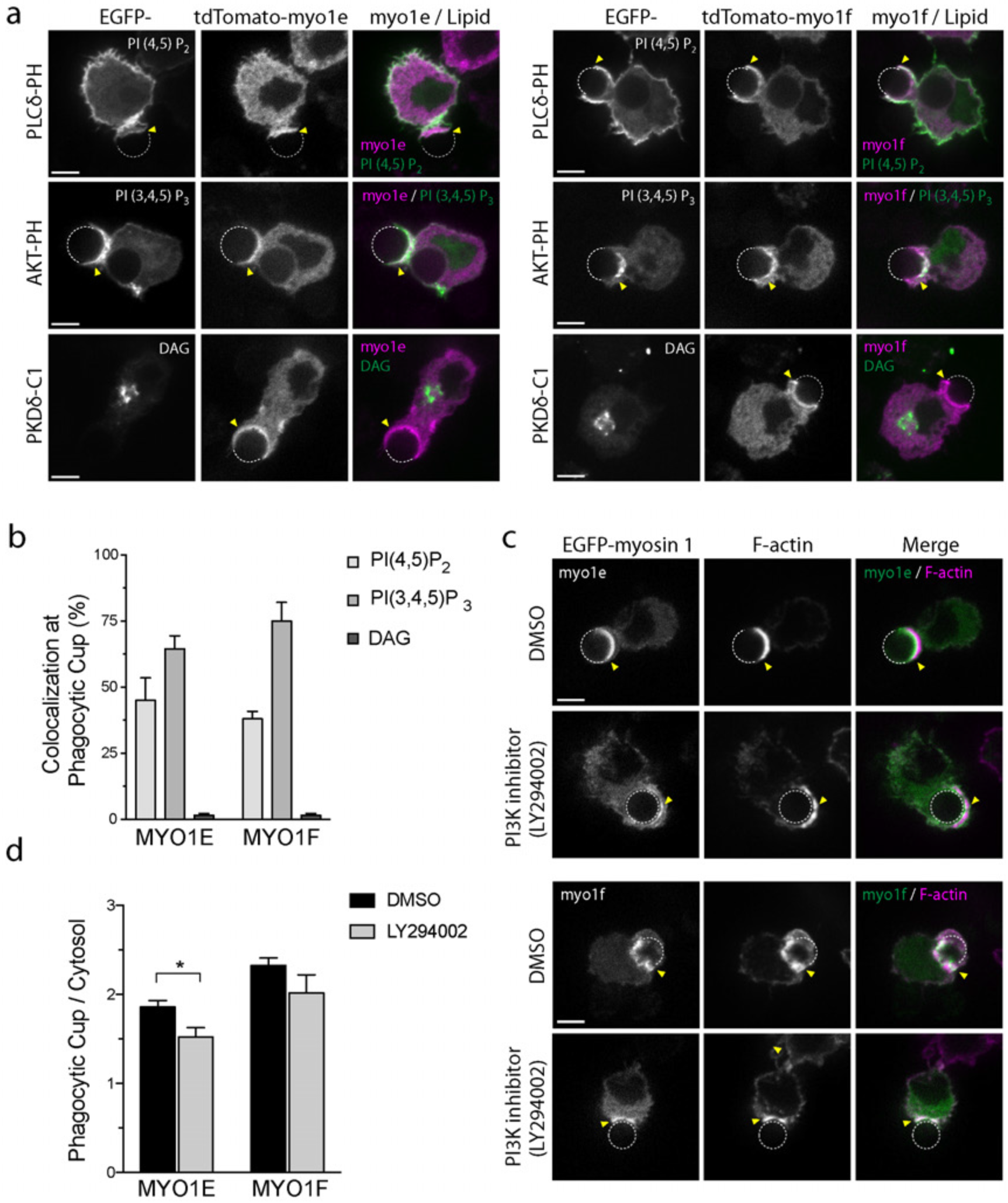
The relationship between myo1e and myo1f localization at the phagocytic cup and phosphoinositides. a) Representative confocal sections of phagocytic cup localization of tdTomato-myo1e/f and EGFP-tagged lipid sensors in RAW macrophages. Yellow arrowheads point to the phagocytic cup, and the bead is outlined by a dotted white line. Scale bar, 5 μm. b) Percentage of cells (mean ± SD) in which myo1e/myo1f colocalized with specific phospholipid markers at the phagocytic cup. EGFP-PKDd-C1 (DAG sensor) was used as a negative control. Data from 2 independent experiments (>50 cups analyzed per pair). c) Myo1e/f localize less robustly to the phagocytic cup when PI3K is inhibited. RAW macrophages transfected with EGFP-myo1e/f were pretreated with DMSO or 50 μM PI3K inhibitor LY294002 for 30 minutes, then challenged to engulf 6 μm IgG-coated beads. Cells were then fixed and stained with phalloidin to label F-actin. Images are representative confocal sections. Yellow arrowheads point to the phagocytic cup, and the bead is outlined by a dotted white line. Scale bar, 5 μm. d) Ratio of myo1e/f enrichment at the phagocytic cup compared to the cytosol (mean ± SEM) in transfected RAW macrophages treated with DMSO or LY294002. Fluorescence intensity levels at the cup and in the cytosol were determined by line scan of single confocal slice (>20 cups analyzed per treatment, *=p<0.05).

**Supplementary Fig. 3:**
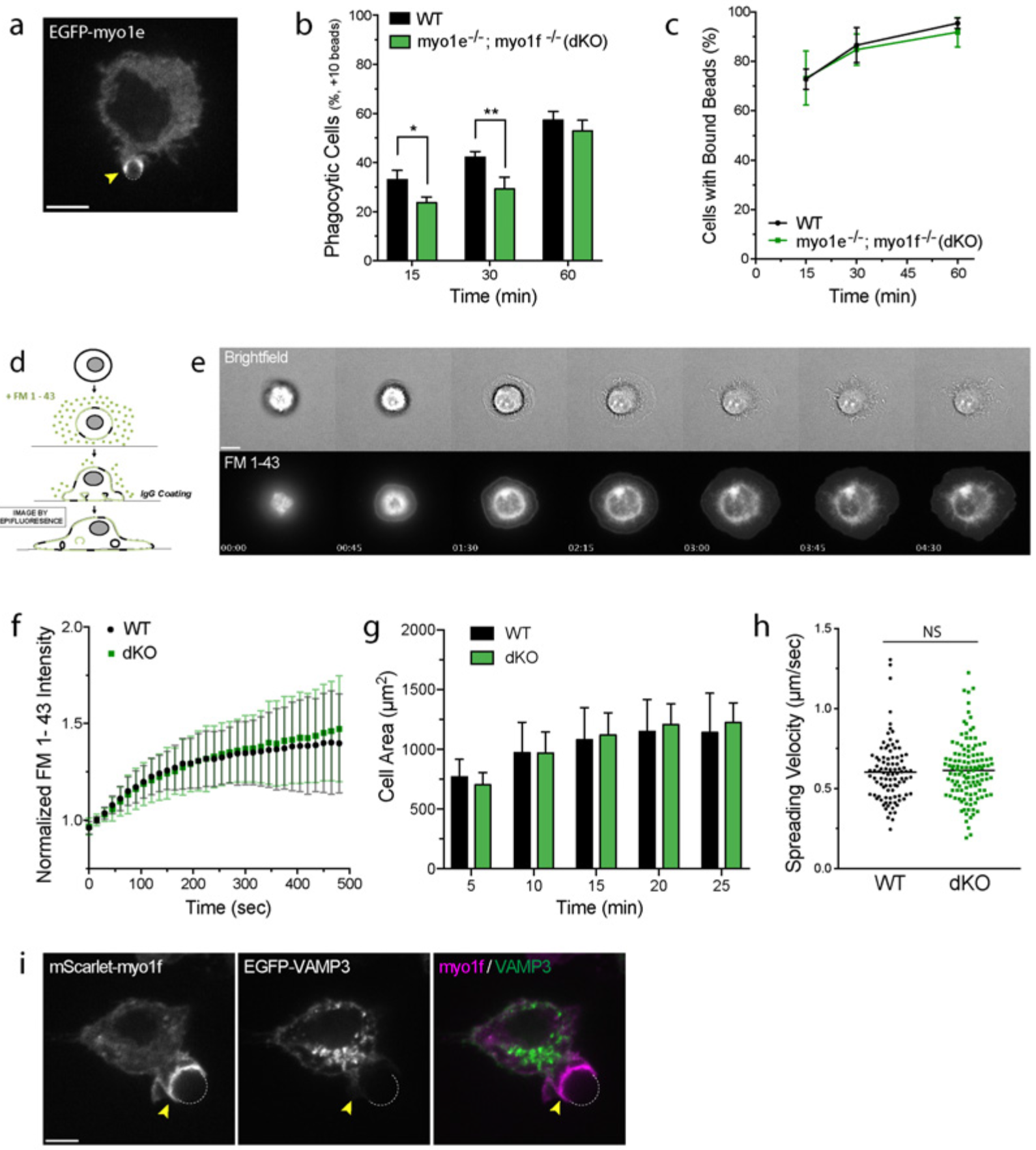
Myo1e/f are not required for focal exocytosis. a) Myo1e is recruited to phagocytic cups formed around small targets. RAW macrophages transfected with EGFP-myo1e were challenged to engulf 2 μm IgG-coated beads. Yellow arrowhead points to the phagocytic cup, and the bead is outlined by a dotted white line. Scale bar, 5 μm. b) Graph of percentage (mean ± SD) of phagocytic cells, defined as those that internalized at least 10 beads. WT and dKO BMDM were challenged with 2 μm IgG-coated latex beads and analyzed at 15, 30, 60 minutes. Data from 3 independent experiments (*= p<0.05 **= p<0.01). c) Percentage of cells (mean ± SEM) that bound at least one small bead during the phagocytosis time course of (b). d) Schematic of the FM 1–43-based membrane quantification experiment. WT and dKO macrophages are serum-starved in suspension and briefly exposed to lipid dye FM 1–43 before being added to a flow chamber to perform frustrated phagocytosis in the presence of excess dye. Fluorescence intensity values are normalized individually to the pre-spread cell. e) Representative time-lapse of WT BMDM performing frustrated phagocytosis and taking up FM 1–43 dye. Cells were imaged using brightfield and wide field fluorescence microscopy. Time stamp at lower left in minutes:seconds. Scale bar, 10 μm. f) Graph (mean ± SD) of normalized FM 1–43 intensity during frustrated phagocytosis. Data from 3 independent experiments (>60 cells per genotype). g) Graph of cell area (mean ± SD) during frustrated phagocytosis. WT and dKO macrophages were serum starved in suspension, then plated on IgG-coated coverslips, and fixed and stained with phalloidin. Data from 3 independent experiments (>100 cells each time point). h) Leading edge velocity of WT and dKO cells during frustrated phagocytosis was measured by kymography. Data from 4 independent experiments (>80 cells per genotype). i) Myo1f and VAMP3 do not colocalize at the phagocytic cup. RAW macrophages co-transfected with EGFP-VAMP3 and mScarlet-myo1f were challenged to engulf 6 μm IgG-coated beads. Yellow arrowhead points to the phagocytic cup, and the bead is outlined by a dotted white line. Scale bar, 5 μm.

**Supplementary Fig. 4:**
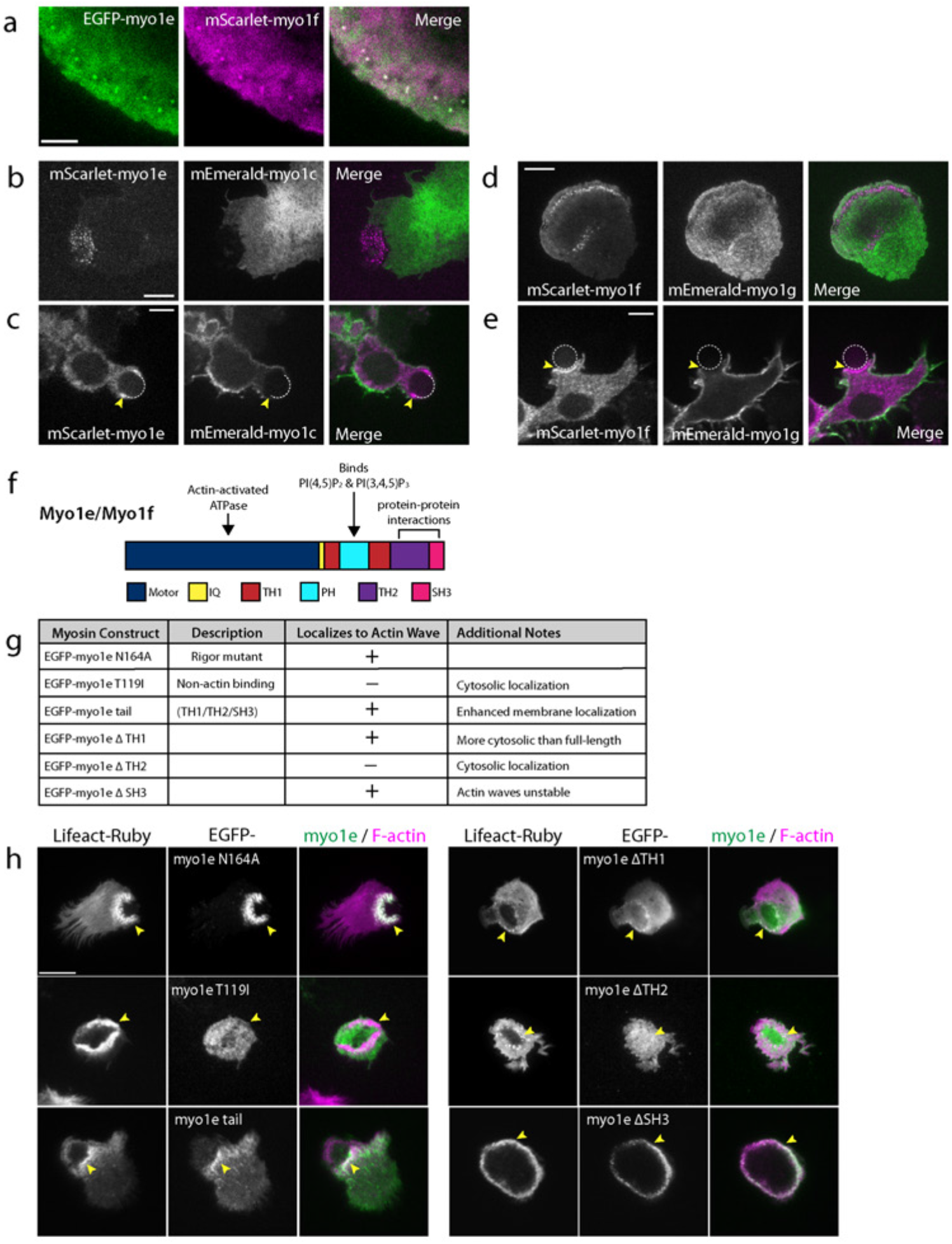
Only long-tailed class 1 myosins (myo1e/f) localize to actin-based adhesions during frustrated phagocytosis. a) Myo1e and myo1f colocalize at punctate adhesions. Spreading edge of RAW macrophage expressing EGFP-myo1e and mScarlet-myo1f conducting frustrated phagocytosis and imaged by TIRFM. Scale bar, 5 μm. b) Myo1c does not colocalize with myo1e at FcR-actin adhesions. Representative TIRFM image of RAW macrophage co-transfected with mScarlet-myo1e and mEmerald-myo1c conducting frustrated phagocytosis. Scale bar, 10 μm. c) Myo1c does not colocalize with myo1e at the phagocytic cup. Representative confocal image of RAW macrophage cotransfected with mScarlet-myo1e and mEmerald-myo1c engulfing a 6 μm IgG-coated bead. Yellow arrowhead points to the phagocytic cup, and the bead is outlined by a dotted white line. Scale bar, 5 μm. d) Myo1g does not colocalize with myo1f at FcR-actin adhesions. Representative TIRFM image of RAW macrophage co-transfected with mScarlet-myo1f and mEmerald-myo1g conducting frustrated phagocytosis. Scale bar, 10 μm. e) Myo1g does not colocalize with myo1f at the phagocytic cup. Representative confocal image of RAW macrophage co-transfected with mScarlet-myo1f and mEmerald-myo1g engulfing a 6 μm IgG-coated bead. Yellow arrowhead points to the phagocytic cup, and the bead is outlined by a dotted white line. Scale bar, 5 μm. f) Schematic of myo1e/f protein domains. N-terminal motor domain produces force on F-actin by ATP hydrolysis. Neck region is defined by a single IQ-motif. Tail homology 1 (TH1) domain contains a putative Pleckstrin Homology (PH) domain to mediate binding to lipids. Tail homology 2 (TH2) and Src Homology 3 (SH3) domain make myo1e/f unique among class 1 myosins and mediate protein-protein interactions. g) Table summarizing myosin colocalization with actin waves for myo1e mutants/deletional constructs during frustrated phagocytosis. Myo1e N164A is a rigor mutant (strongly bound to actin) and T119I is a putative non-actin-binding motor mutant. h) Representative TIRFM images of RAW macrophages, co-expressing EGFP-tagged myo1e mutant/deletional constructs and Lifeact-Ruby, performing frustrated phagocytosis. Yellow arrowheads point to actin waves. Scale bar, 10 μm.

**Supplementary Fig. 5:**
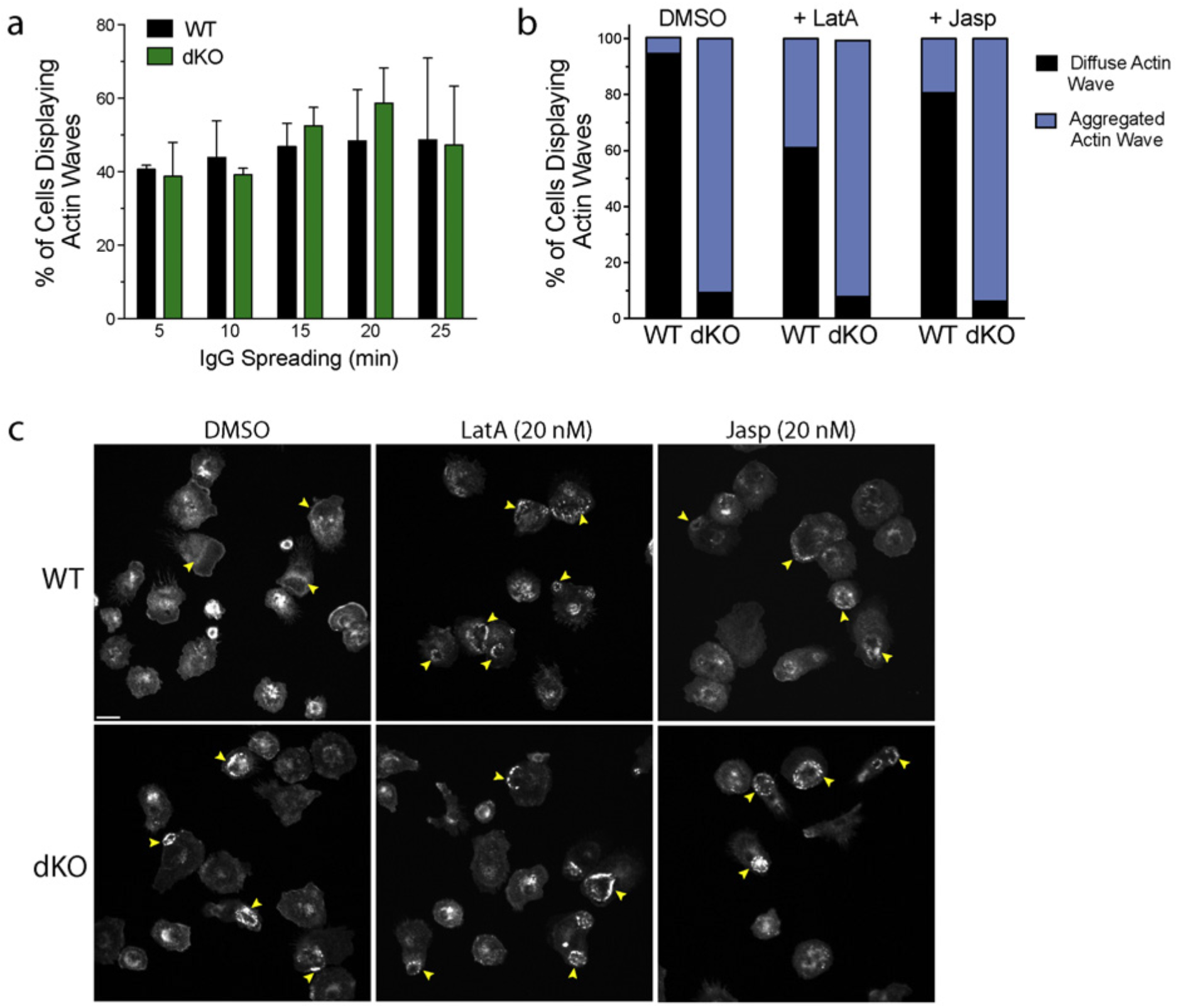
Actin disrupting drugs cannot phenocopy or rescue the defects in actin waves of dKO macrophages. a) Percentage (mean ± SEM) of cells that form actin waves is not different between WT and dKO macrophages over time. WT and dKO BMDM spread on IgG-coated coverslips for specific time points, were then fixed and stained with fluorescently-labeled phalloidin. Data from 3 independent experiments (>100 cells per genotype). b) Treating cells with low doses of Latrunculin A or Jasplakinolide does not rescue actin wave morphology. Graph depicting the mean percentage of cells forming diffuse or aggregated actin waves in WT and dKO BMDM in the presence of 20 nM Latrunculin A or 20 nM Jasplakinolide. Cells were fixed after 10 minutes of frustrated phagocytosis and stained with fluorescently-labeled phalloidin. Data from 2 independent experiments (>70 cells per genotype, judged blindly). c) Representative confocal images of actin waves in WT and dKO BMDM treated with F-actin drugs. Yellow arrows point to actin waves. Scale bar, 20 μm.

**Supplementary Fig. 6:**
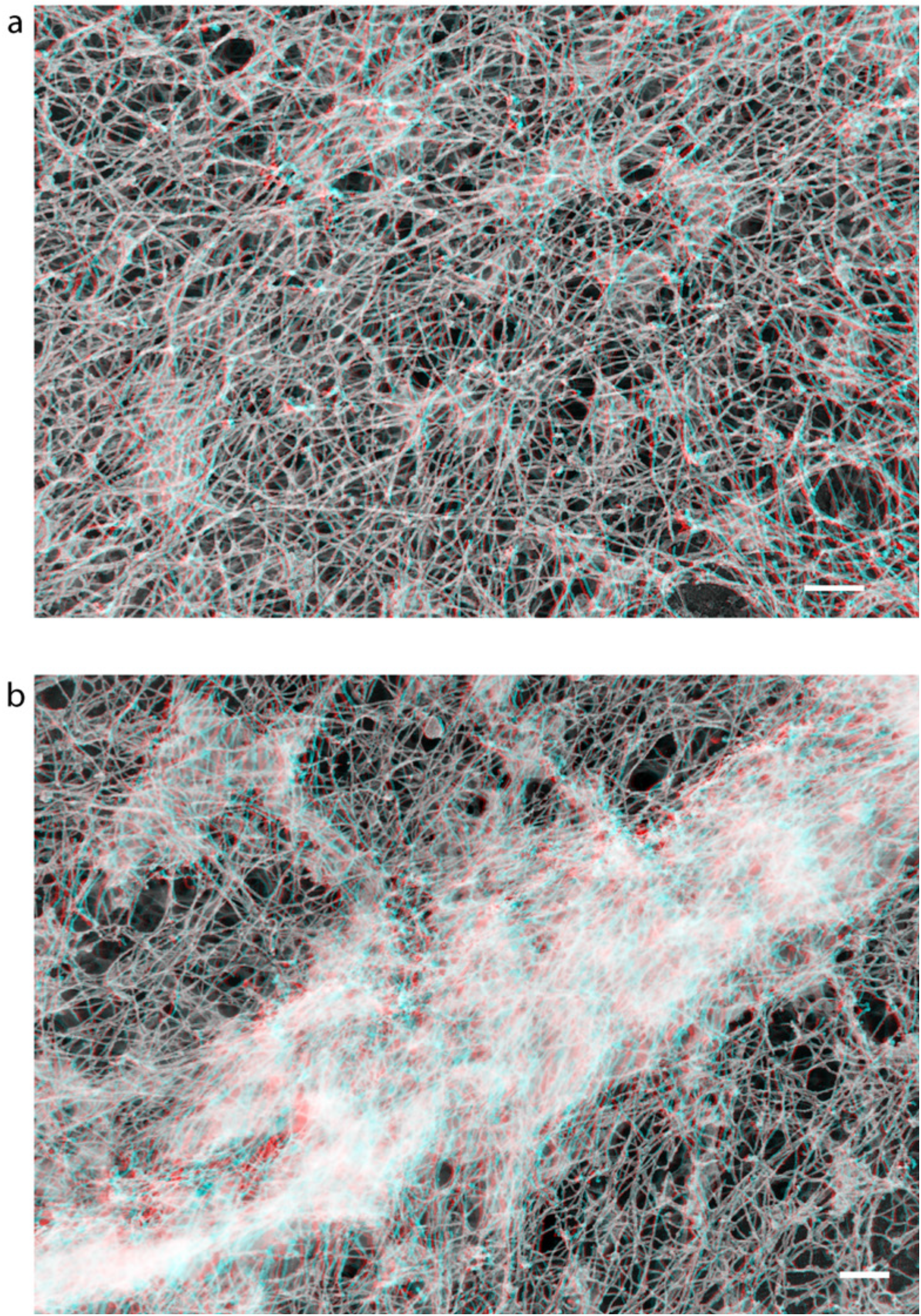
3D platinum replica EM image of actin waves in WT and dKO macrophages. Three-dimensional structure of representative actin wave in WT (a, corresponds to Fig. 5f) and dKO cells (b, corresponds to Fig. 5k). Overlapped stereo pairs of platinum replica electron microscopy images taken at +10 (blue) and –10 (red) degrees of sample tilt. Use 3D view glasses for volume viewing (left eye, blue). Scale bars, 0.2 μm.

## References

1. Aderem, A. & Underhill, D.M. Mechanisms of phagocytosis in macrophages. Annual review of immunology 17, 593–623 (1999).

2. Freeman, S.A. & Grinstein, S. Phagocytosis: receptors, signal integration, and the cytoskeleton. Immunological reviews 262, 193–215 (2014).

3. Braun, V. & Niedergang, F. Linking exocytosis and endocytosis during phagocytosis. Biology of the cell / under the auspices of the European Cell Biology Organization 98, 195–201 (2006).

4. Underhill, D.M. & Goodridge, H.S. Information processing during phagocytosis. Nature reviews. Immunology 12, 492–502 (2012).

5. Garcia-Garcia, E. & Rosales, C. Signal transduction during Fc receptor-mediated phagocytosis. Journal of leukocyte biology 72, 1092–1108 (2002).

6. Lee, C.Y., Herant, M. & Heinrich, V. Target-specific mechanics of phagocytosis: protrusive neutrophil response to zymosan differs from the uptake of antibody-tagged pathogens. Journal of cell science 124, 1106–1114 (2011).

7. Evans, E., Leung, A. & Zhelev, D. Synchrony of cell spreading and contraction force as phagocytes engulf large pathogens. J Cell Biol 122, 1295–1300 (1993).

8. Herant, M., Heinrich, V. & Dembo, M. Mechanics of neutrophil phagocytosis: behavior of the cortical tension. Journal of cell science 118, 1789–1797 (2005).

9. Masters, T.A., Pontes, B., Viasnoff, V., Li, Y. & Gauthier, N.C. Plasma membrane tension orchestrates membrane trafficking, cytoskeletal remodeling, and biochemical signaling during phagocytosis. Proc Natl Acad Sci U S A 110, 11875–11880 (2013).

10. Griffin, F.M., Jr., Griffin, J.A., Leider, J.E. & Silverstein, S.C. Studies on the mechanism of phagocytosis. I. Requirements for circumferential attachment of particle-bound ligands to specific receptors on the macrophage plasma membrane. The Journal of experimental medicine 142, 1263–1282 (1975).

11. Maxeiner, S. et al. Crucial role for the LSP1-myosin1e bimolecular complex in the regulation of Fcgamma receptor-driven phagocytosis. Mol Biol Cell 26, 1652–1664 (2015).

12. Yeung, T. & Grinstein, S. Lipid signaling and the modulation of surface charge during phagocytosis. Immunological reviews 219, 17–36 (2007).

13. Botelho, R.J. et al. Localized biphasic changes in phosphatidylinositol-4,5-bisphosphate at sites of phagocytosis. J Cell Biol 151, 1353–1368 (2000).

14. Marshall, J.G. et al. Restricted accumulation of phosphatidylinositol 3-kinase products in a plasmalemmal subdomain during Fc gamma receptor-mediated phagocytosis. J Cell Biol 153, 1369–1380 (2001).

15. Vieira, O.V. et al. Distinct roles of class I and class III phosphatidylinositol 3-kinases in phagosome formation and maturation. J Cell Biol 155, 19–25 (2001).

16. Feeser, E.A., Ignacio, C.M., Krendel, M. & Ostap, E.M. Myo1e binds anionic phospholipids with high affinity. Biochemistry 49, 9353–9360 (2010).

17. Chen, C.L. & Iijima, M. Myosin I: A new pip (3) effector in chemotaxis and phagocytosis. Communicative & integrative biology 5, 294–296 (2012).

18. Lu, S.M., Grinstein, S. & Fairn, G.D. Quantitative Live-Cell Fluorescence Microscopy During Phagocytosis. Methods in molecular biology 1519, 79–91 (2017).

19. Oancea, E., Teruel, M.N., Quest, A.F. & Meyer, T. Green fluorescent protein (GFP)-tagged cysteine-rich domains from protein kinase C as fluorescent indicators for diacylglycerol signaling in living cells. J Cell Biol 140, 485–498 (1998).

20. Botelho, R.J. et al. Localized diacylglycerol-dependent stimulation of Ras and Rap1 during phagocytosis. J Biol Chem 284, 28522–28532 (2009).

21. Cox, D., Tseng, C.C., Bjekic, G. & Greenberg, S. A requirement for phosphatidylinositol 3-kinase in pseudopod extension. J Biol Chem 274, 1240–1247 (1999).

22. Ninomiya, N. et al. Involvement of phosphatidylinositol 3-kinase in Fc gamma receptor signaling. J Biol Chem 269, 22732–22737 (1994).

23. Krendel, M. et al. Disruption of Myosin 1e promotes podocyte injury. Journal of the American Society of Nephrology : JASN 20, 86–94 (2009).

24. Kim, S.V. et al. Modulation of cell adhesion and motility in the immune system by Myo1f. Science 314, 136–139 (2006).

25. Araki, N., Hatae, T., Furukawa, A. & Swanson, J.A. Phosphoinositide-3-kinase-independent contractile activities associated with Fcgamma-receptor-mediated phagocytosis and macropinocytosis in macrophages. Journal of cell science 116, 247–257 (2003).

26. Swanson, J.A. et al. A contractile activity that closes phagosomes in macrophages. Journal of cell science 112 (Pt 3), 307–316 (1999).

27. Kovari, D.T. et al. Frustrated Phagocytic Spreading of J774A-1 Macrophages Ends in Myosin II-Dependent Contraction. Biophysical journal 111, 2698–2710 (2016).

28. Henson, P.M. Interaction of cells with immune complexes: adherence, release of constituents, and tissue injury. The Journal of experimental medicine 134, 114s–135s (1971).

29. Takemura, R., Stenberg, P.E., Bainton, D.F. & Werb, Z. Rapid redistribution of clathrin onto macrophage plasma membranes in response to Fc receptor-ligand interaction during frustrated phagocytosis. J Cell Biol 102, 55–69 (1986).

30. Labrousse, A.M. et al. Frustrated phagocytosis on micro-patterned immune complexes to characterize lysosome movements in live macrophages. Frontiers in immunology 2, 51 (2011).

31. Masters, T.A., Sheetz, M.P. & Gauthier, N.C. F-actin waves, actin cortex disassembly and focal exocytosis driven by actin-phosphoinositide positive feedback. Cytoskeleton 73, 180–196 (2016).

32. Neuhaus, E.M. & Soldati, T. A myosin I is involved in membrane recycling from early endosomes. J Cell Biol 150, 1013–1026 (2000).

33. Salas-Cortes, L. et al. Myosin Ib modulates the morphology and the protein transport within multi-vesicular sorting endosomes. Journal of cell science 118, 4823–4832 (2005).

34. Schietroma, C. et al. A role for myosin 1e in cortical granule exocytosis in Xenopus oocytes. J Biol Chem 282, 29504–29513 (2007).

35. Schlam, D. et al. Phosphoinositide 3-kinase enables phagocytosis of large particles by terminating actin assembly through Rac/Cdc42 GTPase-activating proteins. Nature communications 6, 8623 (2015).

36. Mohammadi, S. & Isberg, R.R. Cdc42 interacts with the exocyst complex to promote phagocytosis. J Cell Biol 200, 81–93 (2013).

37. Park, H. & Cox, D. Cdc42 regulates Fc gamma receptor-mediated phagocytosis through the activation and phosphorylation of Wiskott-Aldrich syndrome protein (WASP) and neural-WASP. Mol Biol Cell 20, 4500–4508 (2009).

38. Egami, Y., Fukuda, M. & Araki, N. Rab35 regulates phagosome formation through recruitment of ACAP2 in macrophages during FcgammaR-mediated phagocytosis. Journal of cell science 124, 3557–3567 (2011).

39. Braun, V. et al. TI-VAMP/VAMP7 is required for optimal phagocytosis of opsonised particles in macrophages. The EMBO journal 23, 4166–4176 (2004).

40. Weiner, O.D., Marganski, W.A., Wu, L.F., Altschuler, S.J. & Kirschner, M.W. An actin-based wave generator organizes cell motility. PLoS biology 5, e221 (2007).

41. Freeman, S.A. et al. Integrins Form an Expanding Diffusional Barrier that Coordinates Phagocytosis. Cell 164, 128–140 (2016).

42. Lin, J. et al. TIRF imaging of Fc gamma receptor microclusters dynamics and signaling on macrophages during frustrated phagocytosis. BMC Immunol 17, 5 (2016).

43. Dart, A.E., Tollis, S., Bright, M.D., Frankel, G. & Endres, R.G. The motor protein myosin 1G functions in FcgammaR-mediated phagocytosis. Journal of cell science 125, 6020–6029 (2012).

44. Trost, M. et al. The phagosomal proteome in interferon-gamma-activated macrophages. Immunity 30, 143–154 (2009).

45. Bi, J. et al. Effects of FSGS-associated mutations on the stability and function of myosin-1 in fission yeast. Disease models & mechanisms 8, 891–902 (2015).

46. D’Arrigo, C., Candal-Couto, J.J., Greer, M., Veale, D.J. & Woof, J.M. Human neutrophil Fc receptor-mediated adhesion under flow: a hollow fibre model of intravascular arrest. Clinical and experimental immunology 100, 173–179 (1995).

47. Heiple, J.M., Wright, S.D., Allen, N.S. & Silverstein, S.C. Macrophages form circular zones of very close apposition to IgG-coated surfaces. Cell motility and the cytoskeleton 15, 260–270 (1990).

48. Wright, S.D. & Silverstein, S.C. Phagocytosing macrophages exclude proteins from the zones of contact with opsonized targets. Nature 309, 359–361 (1984).

49. Panzer, L. et al. The formins FHOD1 and INF2 regulate inter-and intra-structural contractility of podosomes. Journal of cell science 129, 298–313 (2016).

50. Brandt, D.T. et al. Dia1 and IQGAP1 interact in cell migration and phagocytic cup formation. J Cell Biol 178, 193–200 (2007).

51. May, R.C., Caron, E., Hall, A. & Machesky, L.M. Involvement of the Arp2/3 complex in phagocytosis mediated by FcgammaR or CR3. Nature cell biology 2, 246–248 (2000).

52. Rotty, J.D. et al. Arp2/3 Complex Is Required for Macrophage Integrin Functions but Is Dispensable for FcR Phagocytosis and In Vivo Motility. Developmental cell 42, 498–513 e496 (2017).

53. Pontes, B., Monzo, P. & Gauthier, N.C. Membrane tension: A challenging but universal physical parameter in cell biology. Seminars in cell & developmental biology 71, 30–41 (2017).

54. Diz-Muñoz, A., Weiner, O.D. & Fletcher, D.A. In pursuit of the mechanics that shape cell surfaces. Nature Physics 14, 648–652 (2018).

55. Dai, J. & Sheetz, M.P. Mechanical properties of neuronal growth cone membranes studied by tether formation with laser optical tweezers. Biophysical journal 68, 988–996 (1995).

56. Dai, J. & Sheetz, M.P. Membrane tether formation from blebbing cells. Biophysical journal 77, 3363–3370 (1999).

57. Pontes, B. et al. Membrane tension controls adhesion positioning at the leading edge of cells. J Cell Biol 216, 2959–2977 (2017).

58. Lieber, A.D., Yehudai-Resheff, S., Barnhart, E.L., Theriot, J.A. & Keren, K. Membrane tension in rapidly moving cells is determined by cytoskeletal forces. Current biology : CB 23, 1409–1417 (2013).

59. Brzeska, H., Pridham, K., Chery, G., Titus, M.A. & Korn, E.D. The association of myosin IB with actin waves in dictyostelium requires both the plasma membrane-binding site and actin-binding region in the myosin tail. PloS one 9, e94306 (2014).

60. Gerisch, G. et al. Self-organizing actin waves as planar phagocytic cup structures. Cell adhesion & migration 3, 373–382 (2009).

61. Lynch, T.J. et al. ATPase activities and actin-binding properties of subfragments of Acanthamoeba myosin IA. J Biol Chem 261, 17156–17162 (1986).

62. Jung, G. & Hammer, J.A., 3rd The actin binding site in the tail domain of Dictyostelium myosin IC (myoC) resides within the glycine-and proline-rich sequence (tail homology region 2). FEBS Lett 342, 197–202 (1994).

63. Yu, H.Y. & Bement, W.M. Multiple myosins are required to coordinate actin assembly with coat compression during compensatory endocytosis. Mol Biol Cell 18, 4096–4105 (2007).

64. Kaplan, G. Differences in the mode of phagocytosis with Fc and C3 receptors in macrophages. Scand J Immunol 6, 797–807 (1977).

65. Montesano, R., Mossaz, A., Vassalli, P. & Orci, L. Specialization of the macrophage plasma membrane at sites of interaction with opsonized erythrocytes. J Cell Biol 96, 1227–1233 (1983).

66. Witke, W., Li, W., Kwiatkowski, D.J. & Southwick, F.S. Comparisons of CapG and gelsolin-null macrophages: demonstration of a unique role for CapG in receptor-mediated ruffling, phagocytosis, and vesicle rocketing. J Cell Biol 154, 775–784 (2001).

67. Cheng, J., Grassart, A. & Drubin, D.G. Myosin 1E coordinates actin assembly and cargo trafficking during clathrin-mediated endocytosis. Mol Biol Cell 23, 2891–2904 (2012).

68. Liang, Y., Niederstrasser, H., Edwards, M., Jackson, C.E. & Cooper, J.A. Distinct roles for CARMIL isoforms in cell migration. Mol Biol Cell 20, 5290–5305 (2009).

69. Pernier, J. et al. A new actin depolymerase: a catch bond Myosin 1 motor. bioRxiv (2018).

70. Ikeda, Y. et al. Rac1 switching at the right time and location is essential for Fcgamma receptor-mediated phagosome formation. Journal of cell science 130, 2530–2540 (2017).

71. Tsai, R.K. & Discher, D.E. Inhibition of “self” engulfment through deactivation of myosin-II at the phagocytic synapse between human cells. J Cell Biol 180, 989–1003 (2008).

72. Yamauchi, S., Kawauchi, K. & Sawada, Y. Myosin II-dependent exclusion of CD45 from the site of Fcgamma receptor activation during phagocytosis. FEBS Lett 586, 3229–3235 (2012).

73. Houk, A.R. et al. Membrane tension maintains cell polarity by confining signals to the leading edge during neutrophil migration. Cell 148, 175–188 (2012).

74. Diz-Munoz, A. et al. Membrane Tension Acts Through PLD2 and mTORC2 to Limit Actin Network Assembly During Neutrophil Migration. PLoS biology 14, e1002474 (2016).

75. Batchelder, E.L. et al. Membrane tension regulates motility by controlling lamellipodium organization. Proc Natl Acad Sci U S A 108, 11429–11434 (2011).

76. Herant, M., Lee, C.Y., Dembo, M. & Heinrich, V. Protrusive push versus enveloping embrace: computational model of phagocytosis predicts key regulatory role of cytoskeletal membrane anchors. PLoS computational biology 7, e1001068 (2011).

77. Bi, J. et al. Myosin 1e is a component of the glomerular slit diaphragm complex that regulates actin reorganization during cell-cell contact formation in podocytes. American journal of physiology. Renal physiology 305, F532–544 (2013).

78. Skowron, J.F., Bement, W.M. & Mooseker, M.S. Human brush border myosin-I and myosin-Ic expression in human intestine and Caco-2BBe cells. Cell motility and the cytoskeleton 41, 308–324 (1998).

79. Krendel, M., Osterweil, E.K. & Mooseker, M.S. Myosin 1E interacts with synaptojanin-1 and dynamin and is involved in endocytosis. FEBS Lett 581, 644–650 (2007).

80. Fitzer-Attas, C.J. et al. Fcgamma receptor-mediated phagocytosis in macrophages lacking the Src family tyrosine kinases Hck, Fgr, and Lyn. The Journal of experimental medicine 191, 669–682 (2000).

81. Greuber, E.K. & Pendergast, A.M. Abl family kinases regulate FcgammaR-mediated phagocytosis in murine macrophages. Journal of immunology 189, 5382–5392 (2012).

82. Svitkina, T.M. & Borisy, G.G. Correlative light and electron microscopy of the cytoskeleton of cultured cells. Methods in enzymology 298, 570–592 (1998).

83. Svitkina, T. Electron microscopic analysis of the leading edge in migrating cells. Methods Cell Biol 79, 295–319 (2007).

84. Svitkina, T. Imaging cytoskeleton components by electron microscopy. Methods in molecular biology 586, 187–206 (2009).

85. Oakes, P.W. et al. Lamellipodium is a myosin-independent mechanosensor. Proc Natl Acad Sci U S A 115, 2646–2651 (2018).

86. Sabass, B., Gardel, M.L., Waterman, C.M. & Schwarz, U.S. High resolution traction force microscopy based on experimental and computational advances. Biophysical journal 94, 207–220 (2008).

87. Oakes, P.W., Banerjee, S., Marchetti, M.C. & Gardel, M.L. Geometry regulates traction stresses in adherent cells. Biophysical journal 107, 825–833 (2014).

88. Li, Q.S., Lee, G.Y., Ong, C.N. & Lim, C.T. AFM indentation study of breast cancer cells. Biochemical and biophysical research communications 374, 609–613 (2008).

89. Wong, W.P. & Halvorsen, K. The effect of integration time on fluctuation measurements: calibrating an optical trap in the presence of motion blur. Optics express 14, 12517–12531 (2006).

90. Schindelin, J.; Arganda-Carreras, I. & Frise, E. et al. Fiji: an open-source platform for biological-image analysis. Nature methods 9: 676–682 (2012).

91. R Core Team. R: A language and environment for statistical computing. R Foundation for Statistical Computing, Vienna, Austria. URL https://www.Rproject.org/. (2018)

92. Wickham, H.. ggplot2: Elegant Graphics for Data Analysis. Springer-Verlag New York (2016).

